# Structural organization of the C1a-e-c supercomplex within the ciliary central apparatus

**DOI:** 10.1101/773416

**Authors:** Gang Fu, Lei Zhao, Erin Dymek, Yuqing Hou, Kangkang Song, Nhan Phan, Zhiguo Shang, Elizabeth F. Smith, George B. Witman, Daniela Nicastro

## Abstract

Nearly all motile cilia contain a central apparatus (CA) composed of two connected singlet-microtubules with attached projections that play crucial roles in regulating ciliary motility. Defects in CA assembly usually result in motility-impaired or paralyzed cilia, which in humans causes disease. Despite their importance, the protein composition and functions of the CA projections are largely unknown. Here, we integrated biochemical and genetic approaches with cryo-electron tomography to compare the CA of wild type *Chlamydomonas* with CA mutants. We identified a large (>2 MDa) complex, the C1a-e-c supercomplex, that requires the PF16 protein for assembly and contains the CA components FAP76, FAP81, FAP92, and FAP216. We localized these subunits within the supercomplex using nanogold-labeling and show that loss of any one of them results in impaired ciliary motility. These data provide insight into the subunit organization and three-dimensional (3D) structure of the CA, which is a prerequisite for understanding the molecular mechanisms by which the CA regulates ciliary beating.

**Summary:** Fu et al. use a wild-type vs. mutant comparison and cryo-electron tomography of *Chlamydomonas* flagella to identify central apparatus (CA) subunits and visualize their location in the native 3D CA structure. The study provides a better understanding of the CA and how it regulates ciliary motility.

## Introduction

Cilia and flagella are highly conserved organelles in eukaryotes. They have roles in cell motility, generating fluid flow, and sensing extracellular cues. Defects in ciliary assembly or function cause a wide range of human diseases, collectively termed ciliopathies (Afzelius, 2004; Fliegauf et al., 2007). The “9+2” axonemal core structure of motile cilia consists of nine outer doublet microtubules (DMTs) surrounding two singlet microtubules (C1 and C2) that form the central apparatus (CA), or central pair complex. Attached to this axonemal microtubule scaffold are hundreds of proteins (Pazour et al., 2005), including the inner and outer arm dynein motors, and regulatory complexes forming part of the signal transduction pathway(s) that coordinate dynein activity to generate ciliary motility (Summers and Gibbons, 1971; Sale and Satir, 1977; Lin and Nicastro, 2018; Witman et al., 1978; Smith and Sale, 1992; Piperno et al., 1994; Smith and Lefebvre, 1997a; Porter and Sale, 2000; Smith, 2002; Mitchell, 2004; Nicastro et al., 2006; Dymek and Smith, 2007; Wirschell et al., 2007; Bower et al., 2009; Heuser et al., 2009; b; a; Yamamoto et al., 2013; Loreng and Smith, 2017; Fu et al., 2018; Kubo et al., 2018).

The CA is the largest known ciliary regulatory complex. Early structural analyzes described the CA as an asymmetric assembly with seven C1- and C2-projections, but our previous cryo-electron tomography (cryo-ET) study of the wild type *Chlamydomonas* CA revealed at least eleven projections that have 16–32 nm periodicities along the ciliary length, and form connections between C1 and C2, as well as to the radial spoke heads (Witman et al., 1978; Dutcher, 1984; Mitchell and Sale, 1999; Mitchell, 2003; Mitchell and Smith, 2009; Carbajal-González et al., 2013; Loreng and Smith, 2017). Mutations of CA components often result in impaired or paralyzed cilia (Witman et al., 1978; Dutcher, 1984; Smith and Lefebvre, 1996; Smith and Lefebvre, 1997b; Smith and Yang, 2004). Deficiency of CA proteins can cause mammalian ciliopathies, including primary ciliary dyskinesia (PCD) (Teves et al., 2016; Horani and Ferkol, 2018). Mice deficient in either c*fap54* or *pcdp1*, which encode conserved C1d proteins, show typical PCD symptoms, including ineffective mucus clearance, and/or male infertility, and hydrocephalus (Lee et al., 2008; DiPetrillo and Smith, 2010; McKenzie et al., 2015). Similarly, mutations of the C2b protein Hydin result in hydrocephalus caused by the loss of cilia-generated fluid flow in the brain ventricles (Lechtreck et al., 2008).

Despite many biochemical and structural studies of the CA (Witman et al., 1978; Dutcher, 1984; Smith and Lefebvre, 1997a; Mitchell, 2003; Mitchell and Smith, 2009; Loreng and Smith, 2017), the protein composition, 3D organization, and functional mechanism(s) of the CA in ciliary motility are not fully understood. Our recent mass spectrometry (MS) study compared the proteomes of *Chlamydomonas* wild-type and mutant axonemes, and identified 44 new candidate CA proteins assigned to the C1 or the C2 microtubule (Zhao et al., 2019). However, questions about the organization, assembly, and function of the CA and its projections remain, making the CA the structurally and functionally least understood axonemal complex to date.

Here we combined biochemical, genetic, and structural analyses to investigate the protein composition and molecular organization of a group of interconnected CA projections, here termed the C1a-e-c supercomplex, in wild-type and CA mutants of *Chlamydomonas*. Sucrose gradient sedimentation and MS revealed that several CA proteins identified in this study, FAP76, FAP81, FAP92 and FAP216, are associated with the protein PF16, previously assigned to the C1 microtubule but not to a specific projection. *Chlamydomonas* mutants that lacked any of these proteins showed impaired motility. Structural comparisons of flagella from wild-type, these mutants, and tagged rescue strains revealed the precise locations of PF16, FAP76, FAP81, FAP92, and FAP216 within the C1a-e-c supercomplex. Our data show that stable assembly of this supercomplex and its interaction with the neighboring C1d projection are required for the proper regulation of ciliary motility.

## Results

### An improved wild-type CA flagella structure

Cilia were isolated from *Chlamydomonas* cells, demembranated and frozen rapidly for cryo-ET imaging and subtomogram averaging of the DMT and CA repeats. Our previous cryo-ET study of the wild-type CA structure of *Chlamydomonas* flagella achieved 3.5 nm resolution (FSC 0.5 criterion) (Fig. 1 A, C-E) (Carbajal-González et al., 2013). Here, we improved the resolution of the CA structure to 2.3 nm (Fig. 1 B, C, F and G) by applying advanced hardware and software. For example, tilt series were recorded with multiple frames per image (to correct for beam-induced sample-motion; Brilot et al., 2012) on a direct electron detector (Cheng et al., 2015), using a Volta-Phase-Plate (to improve image contrast close to focus; Danev et al., 2014) and a dose-symmetric tilting scheme (to reduce radiation damage of the sample in images recorded at low tilt angles; Hagen et al., 2016). This improves the resolution of the subtomogram averages of axonemes, shown by averages of DMT-associated structures with up to 1.8 nm resolution (FSC 0.5 criterion; Lin et al., 2019).

**Figure 1.**
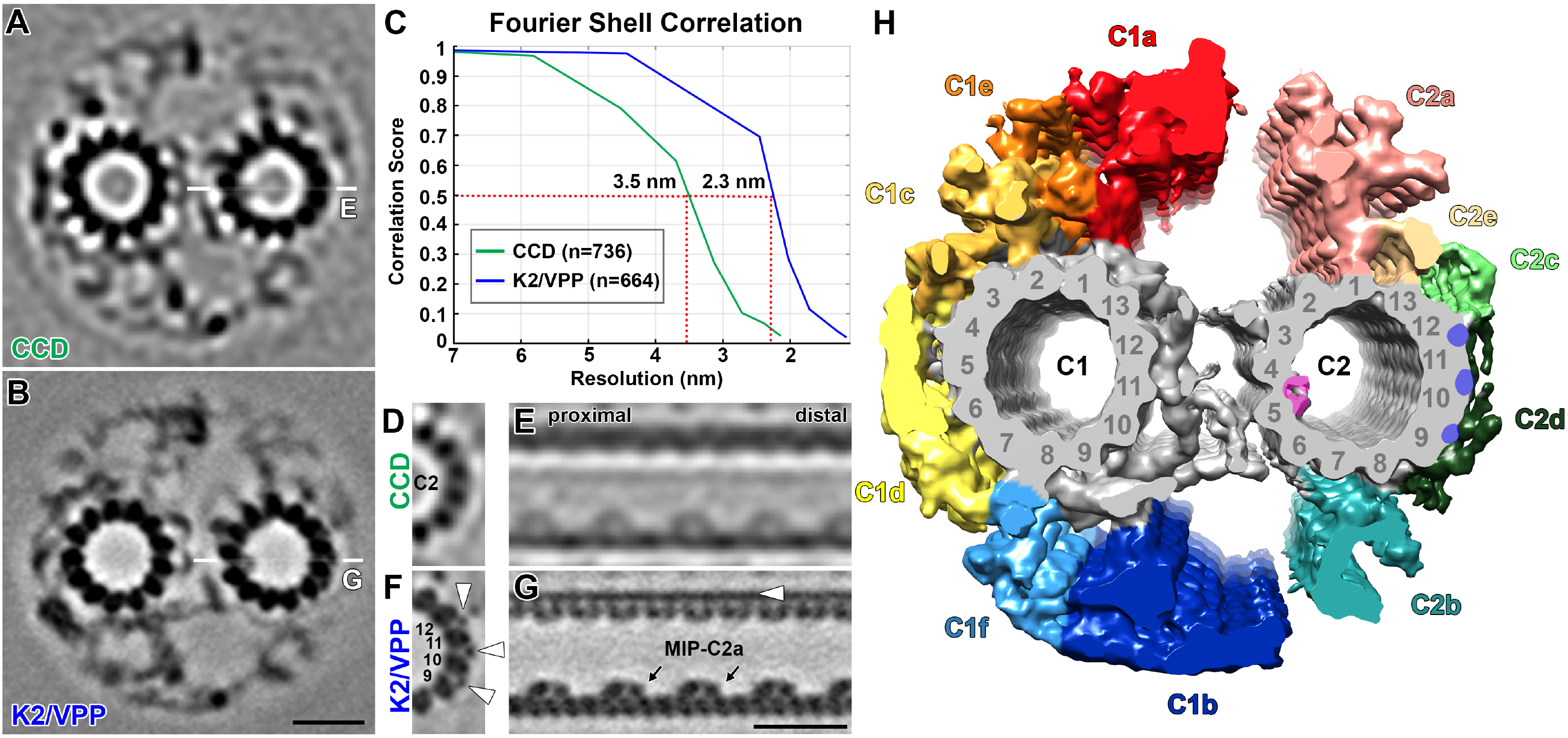
Improved resolution of the averaged CA structure. **(A** and **B)** Tomographic slices of the averaged 32 nm repeat of the *Chlamydomonas* wild-type CA viewed in cross-section to compare data recorded either with a charge-coupled device (CCD) camera (A) or with a direct electron detector and Volta Phase Plate (K2/VPP) (B). Thin white lines in A and B indicate the locations of the slices shown in E and G, respectively. **(C)** Fourier shell correlation curves of the CA averages show that the resolution is improved from 3.5 nm for CCD data to 2.3 nm for K2/VPP data (0.5 criterion). **(D-G)** Tomographic slices to compare the C2 microtubule-associated structures between the CCD (D and E) and the K2/VPP data (F and G) in cross-sectional (D and F) and longitudinal views (E and G). Note the clear visualizations of (i) filamentous, microtubule-associated proteins between protofilaments 9-12 (white arrowheads in F and G), (ii) two distinct domains of the microtubule inner protein (MIP) C2a (black arrows in G), and (iii) the tubulin lattice of the microtubule wall (G) in the K2/VPP data, which were not observed or were blurred in the CCD data. **(H)** Isosurface rendering shows the averaged *Chlamydomonas* wild-type CA (K2/VPP data) in cross-sectional view. Naming and coloring of CA projections adopted from Mitchell and Sale, 1999; Carbajal-González et al., 2013. CA protofilaments were not previously numbered; here we assigned protofilament #1 of C1 and C2 to where the C1a and C2a projections attach, respectively. In all figures, cross-sections are viewed from proximal (i.e., cell body) to the ciliary tip, and in longitudinal views the proximal side is on the right. Scale bar in B, 20 nm (valid for A, B); in G, 20 nm (valid for D-G).

The improved 3D structure of the *Chlamydomonas* wild-type CA reveals details of the CA microtubules, the bridge between them, and the various CA projections (Fig. 1 F-H and Video 1). Previously unresolved details include: (a) Three filamentous structures, with a <2 nm diameter, between protofilaments 9-12 of the C2 microtubule (compare Fig. 1 D and F); (b) the Microtubule Inner Protein, MIP-C2a, which forms an arch-shaped structure on the inside of the C2 wall (Carbajal-González et al., 2013) (Fig. 1 E), consists of two distinct sub-structures that are bound to alternating tubulin subunits of protofilament 5 (Fig. 1 G); (c) the C2a projection, previously shown to have a 8 nm periodicity (Carbajal-González et al., 2013), instead has a 16 nm periodicity and adopts two conformational states (Fig. S1 A-E); (d) two connections between the C1a and C1e projections, and (e) a peripheral density at the C1c projection (Fig. S1 F and G).

### A PF16-dependent complex

A recent CA proteomics study identified several candidate C1-proteins that are reduce or absent from isolated axonemes of the *Chlamydomonas* “paralyzed flagella” mutant *pf16* (Zhao et al., 2019), which lacks the entire C1 microtubule and its associated projections (Dutcher, 1984; Smith and Lefebvre, 1996). To identify proteins that are closely associated with the PF16 subunit, and to visualize their locations within distinct C1-projections using cryo-ET, we narrowed down the list of C1-candidate proteins to those that co-sediment with PF16. Rather than comparing the co-sedimentation profiles of the full axonemal proteomes between wild type and *pf16*, we limited the proteome of interest by comparing the *pf28* mutant, which lacks the large ODA complexes, but contains the CA (Mitchell and Rosenbaum, 1985; Kamiya, 1988), with the double-mutant *pf28;pf16*, and by extracting proteins from isolated axonemes using high potassium iodide (KI) concentration, which eliminates the DMTs and proteins strongly associated with the DMTs from the sample, before separating the KI-extracts on sucrose gradients under native conditions.

We probed the sucrose gradient fractions of the *pf28* extract with anti-PF16 antibody, analyzed the positive fractions with MS, and subtracted those proteins that were also found in comparable fractions of the C1-less double-mutant *pf28;pf16* (Table S1). Cross-referencing the proteins of this difference set with previous studies identified 16 potential PF16-associated proteins (Table 1): Five proteins (PF6, FAP101, FAP114, FAP119, and FAP227) were previously assigned with the C1a projection and two with the C1d projection (FAP221 and FAP54), whereas PP1c (Yang et al., 2000) and eight recently identified candidate C1-proteins (Zhao et al., 2019) had not yet assigned to specific C1 projections. Of these nine unassigned proteins FAP76, FAP81, FAP92, and FAP216, were large enough to be suitable targets for cryo-ET imaging. *Chlamydomonas* mutants corresponding to *FAP76* (*fap76-1* and *fap76-2*), *FAP81*, *FAP92*, and *FAP216*, were purchased from the *Chlamydomonas* Library Project (CLiP) (Li et al., 2016). Mutations were verified by PCR (Fig. S2 A), MS (Table 1 and Table S2), and rescue the mutant phenotypes (motility and structure) by transformation with the corresponding wild-type genes. We also generated a *fap76-1;fap81* double mutant.

**Figure 2.**
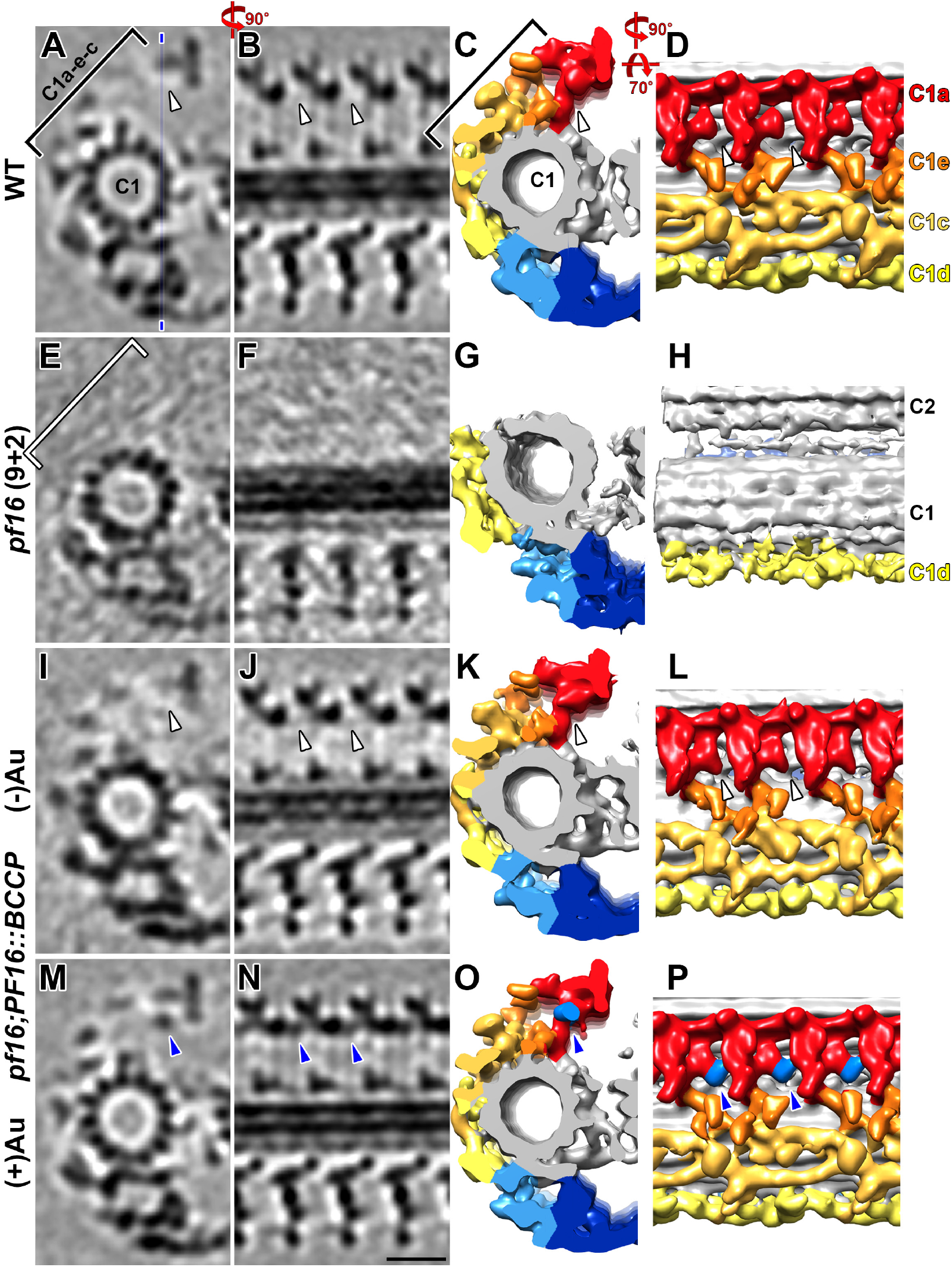
CA projections C1a, e, and c are lost in the *pf16* mutant, and the PF16 C-terminus locates to the C1a projection. **(A-P)** Tomographic slices (columns 1 and 2) and isosurface renderings (columns 3 and 4) of the averaged CA repeats of wild type (A-D), “9+2” *pf16* (E-F), and of the tagged rescue *pf16;PF16::BCCP* without adding nanogold (I-L, control) and after treatment with streptavidin gold (M-P) in cross-sectional (columns 1 and 3) and longitudinal views (columns 2 and 4). The thin blue line in (A) indicates the location of the slices shown in column 2. The C1a, e and c projections (indicated by black brackets in A and C) were missing in the *pf16* mutant (white bracket in E). When the C-terminus of PF16 was tagged with BCCP, the additional density of the BCCP-streptavidin-gold label was detected in the C1a projection (blue arrowheads in M-P); this label density was not observed in wild type (white arrowheads in A-D) or the control samples (white arrowheads in I-L). Scale bar in N, 20 nm (valid for all EM images).

### PF16, a C1a projection subunit, is required for C1a-e-c complex assembly

A previous conventional EM study showed that in the *pf16* mutant the C1 microtubule and its projections were unstable upon isolation of axonemes, i.e. only 8% of *pf16* axonemes contained the C1 and C2 microtubules (“9+2” axonemes), whereas most of the intact *pf16* flagella had both CA microtubules, as did flagella and isolated axonemes from wild-type *Chlamydomonas* (Dutcher et al., 1984; Mitchell and Sale, 1999). However, the resolution in previous studies was insufficient to visualize the extent of missing C1-projection(s) in *pf16* axonemes containing the C1 microtubule. Therefore, to characterize the CA structure that requires PF16 protein for its stable assembly, we performed cryo-ET and subtomogram averaging of those *pf16* CAs that still contained a C1 microtubule (“9+2”) and compared the averages to the wild-type CA structure. Of 39 cryo-tomograms of *pf16* axonemes only 10% had both microtubules (“9+2”), whereas the remaining axonemes lacked either the C1 microtubule (“9+1”, 51%) or the entire CA (“9+0”, 39%), consistent with previous studies (Dutcher, 1984; Mitchell and Sale, 1999), but the subtomogram averages revealed that in the *pf16* “9+2” axonemes the C1d, C1f, and C1b projections remained structurally intact and only the C1a, C1e, and C1c projections, with an estimated total mass of ∼2.0 MDa, were absent (Fig. 2 E-H)^1^. Thus, the C1a-e-c projections require PF16 for stable assembly onto the C1 microtubule, and they form a protein interaction network suggestive of a “C1a-e-c supercomplex” within the ciliary CA.

Although the structural defect in the *pf16* mutant was informative about the extent of the PF16-dependent protein network, it did not reveal the location of the PF16 protein within the C1a-e-c supercomplex. To determine the location of PF16, we used the clonable biotin-carboxyl-carrier-protein (BCCP) tag developed for visualizing gene products by cryo-ET and subtomogram averaging (Oda and Kikkawa, 2013; Song et al., 2015). We rescued the *pf16* mutant with wild-type *PF16* C-terminally tagged with BCCP. Axonemes of the rescued strain were isolated, and the biotinylated tag was enhanced with streptavidin-nanogold, which is visible as additional cryo-EM density in comparison to the wild-type structure (Song et al., 2015; Fu et al., 2018). Tomograms of the *pf16;PF16::BCCP* rescue showed only “9+2” axonemes and all CA projections were restored to the wild-type architecture with and without (control) addition of streptavidin-gold (Fig. 2 I-P), confirming that stable assembly of the C1 microtubule and C1a-e-c supercomplex requires PF16 protein. In contrast to the wild-type (Fig. 2 A-D) and control axonemes without streptavidin-gold (Fig. 2 I-L), the CA average of *pf16;PF16::BCCP* with streptavidin-gold revealed an additional cryo-EM density in the periphery of the C1a projection (blue arrowheads in Fig. 2 M-P, Video 2), indicating the location of the PF16 C-terminus.

We were unable to rescue the *pf16* mutant with N-terminally tagged PF16, possibly because the PF16 N-terminus is required for protein-protein interactions and assembly of the C1a-e-c supercomplex, or the N-terminal BCCP tag (9 kDa) disrupts PF16 folding or its transport into the axoneme.

### C1a-e-c supercomplex mutants have defective swimming and photoshock response

CA mutants often have paralyzed flagella (see *pf16* in Fig. 3 A). In contrast, the *fap76-1*, *fap81*, *fap92,* and *fap216* cells swim, but with slower velocity and a curving swimming path, caused by lack of synchronization between the two flagella (Video 3), rather than the straight or a loose helical paths of wild-type cells (Fig. 3 A and B, Video 3). Among the four mutants, *fap92* has the mildest motility impairment (70% wild-type swimming speed and only a slight curving path). Cells from *fap76-1*, *fap81*, *fap216* and the double-mutant *fap76-1;fap81* have similarly severe motility defects with slower velocity (∼50% of wild type) and clearly curving swimming paths (Fig. 3 A and B). Flagella asynchrony has also been reported for a *Chlamydomonas fap74* RNAi mutant that lacks the C1d projection (DiPetrillo and Smith, 2011).

**Figure 3.**
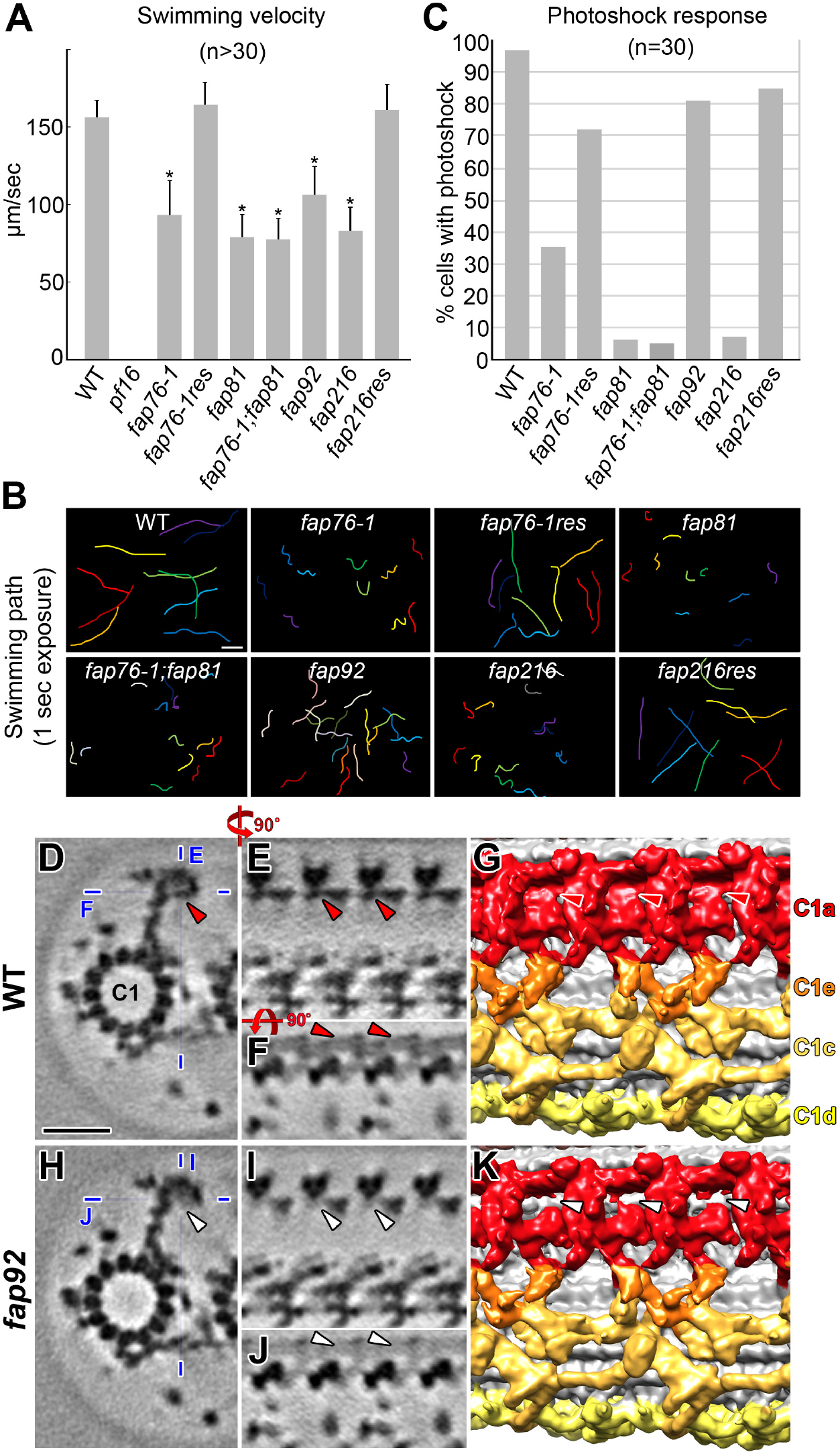
Motility defects of C1a-e-c mutants, and FAP92 is a C1a protein. **(A)** Average swimming velocities of wild-type, mutant cells, and rescue strains *fap76-1;BCCP::FAP76* (*fap76-1res*) and *fap216;BCCP::FAP216* (*fap216res*). In the sequence as shown in the histogram n=30, 30, 30, 30, 30, 30, 32, 40, 34. Asterisks (*) indicate a significant difference (student test, *p*<0.01) compared to wild type. Error bars indicate ± s.e.m. **(B)** Each colored line represents the swimming path of one cell recorded for 1 s. The swimming paths of mutants are curving. **(C)** The percentage of cells displaying photoshock response, i.e. that switched from forward swimming to backwards swimming upon light stimulus, compared to wild type. (**D**-**K**) Comparisons of tomographic slices (D-F, H-J) and isosurface renderings (G and K) between the averaged CA repeats of wild-type (D-G) and *fap92* axonemes (H-K) viewed in cross-sectional (D and H), longitudinal (E, G, I and K) and top-down (F and J) orientations, showing a sheet-like density that is present in the wild-type C1a projection (red arrowheads in D-G) but missing in *fap92* (white arrowheads in H-K). Thin blue lines in D and H indicate the location of the tomographic slices shown in (E and F) and (I and J), respectively. Scale bar in B (WT), 50 µm (valid for all images in B); in D, 20 nm (valid for D-F and H-J).

Previous studies have shown that C1d-less mutants lack a proper photoshock response (DiPetrillo and Smith, 2011; Brown et al., 2012). Our results show that the photoshock response was also impaired in the C1a-e-c supercomplex CLiP mutants, i.e. only 35%, 6%, 7%, and 5% of *fap76-1*, *fap81*, *fap216* and double-mutant *fap76-1;fap81* cells, respectively, exhibited photoshock, whereas *fap92* had a milder photoshock defect (81%; Fig. 3 C). The wild-type swimming speed and straight swimming path were completely restored in the rescue strains *fap76-1;BCCP::FAP76* and *fap216;BCCP::FAP216* (Fig. 3 A and B); however, the photoshock response of *fap76-1;BCCP::FAP76* was still slightly impaired (72%; Fig. 3 C). The similar phenotypes of the C1a-e-c and previously reported C1d mutants suggest that these CA projections might functionally interact to regulate ciliary motility.

### Subunit localization within specific CA projections

The impaired cell motility, and the perturbed proteome of the C1a-e-c supercomplex mutants, suggest defects in their flagellar structure. Therefore, we used cryo-ET and subtomogram averaging to compare the CLiP mutant CA and control DMT structures with wild-type axonemes. None of the mutant averages showed DMT-associated defects (Fig. S3 A-G), but all of them revealed structural defects in the CA with a loss of mass ranging from 80 kDa to 1.2 MDa.

#### FAP92 is a C1a protein that connects neighboring C1a projections

Compared to wild type, the *fap92* CA is missing a sheet-like C1a density (compare Fig. 3 D-G with H-K; Video 2). In the wild-type CA, this sheet-like structure has a 16 nm periodicity along the axonemal length and contributes to the connection between neighboring C1a projections in the periphery of the CA (Fig. 3 G). The MS analyses identified similar numbers of unique peptides throughout the predicted amino acid sequence of FAP92 in both wild-type and *fap92* axonemes suggesting that FAP92 is present in isolated *fap92* axonemes (Table 1). However, sequencing of the genomic DNA showed that the cassette was inserted into the 13th exon of *FAP92* in the mutant genome (Fig. S2 A), which was accompanied by the deletion of three nucleotides (ACC) immediately upstream of the insertion site in exon 13 (unpublished results). The remainder of exon 13 was intact downstream of the cassette. RT-PCR confirmed that the coding region of *FAP92* was disrupted, with a fragment corresponding to exons 12-15 being amplified from wild-type but not from *fap92* cDNA (Fig. S2 B).

The mutant FAP92 protein structure is likely altered, causing it to be partially disordered or positionally flexible and thus not visible in the axonemal averages, and/or precluded from forming stable interactions with FAP413, which was significantly reduced in *fap92* axonemes (Table S2). The remaining C1a structures are not destabilized in the *fap92* mutant (e.g. the classification analyses did not reveal structural flexibility of the C1a projections), and neighboring C1a projections are still connected through additional linking structures (Fig. 3 K), even though *fap92* cells swim slowly, indicative of impaired flagellar motility. Thus, FAP92 probably has a role in regulating ciliary beating, as opposed to having a major scaffolding function.

#### FAP76 is an C1c protein that interacts with the C1e and C1d projections

Cryo-ET and subtomogram averaging of the *fap76-1* mutant revealed that the outer part of the C1c projection was missing in the mutant CA compared to wild type (compare Fig. 4 A-D with E-H). The FAP76-dependent density has a triskelion-like shape with 32 nm periodicity along the C1 microtubule (magenta in Fig. 4 I). MS analysis of *fap76-1* axonemes revealed that FAP76 is significantly reduced (Table 1), and the estimated size of the triskelion-shaped density of ∼190 kDa suggests it contains a single copy of FAP76 (170 kDa). FAP76 forms at least three connections with neighboring structures involving three different CA projections: #1 is the interaction between FAP76 and other C1c densities near the interface between the C1c and C1e projections (Fig. 4 I; Video 2); #2 is the attachment of FAP76 to a rod-shaped C1d density, and #3 links two C1d projections (Fig. 4 J; Video 2).

**Figure 4.**
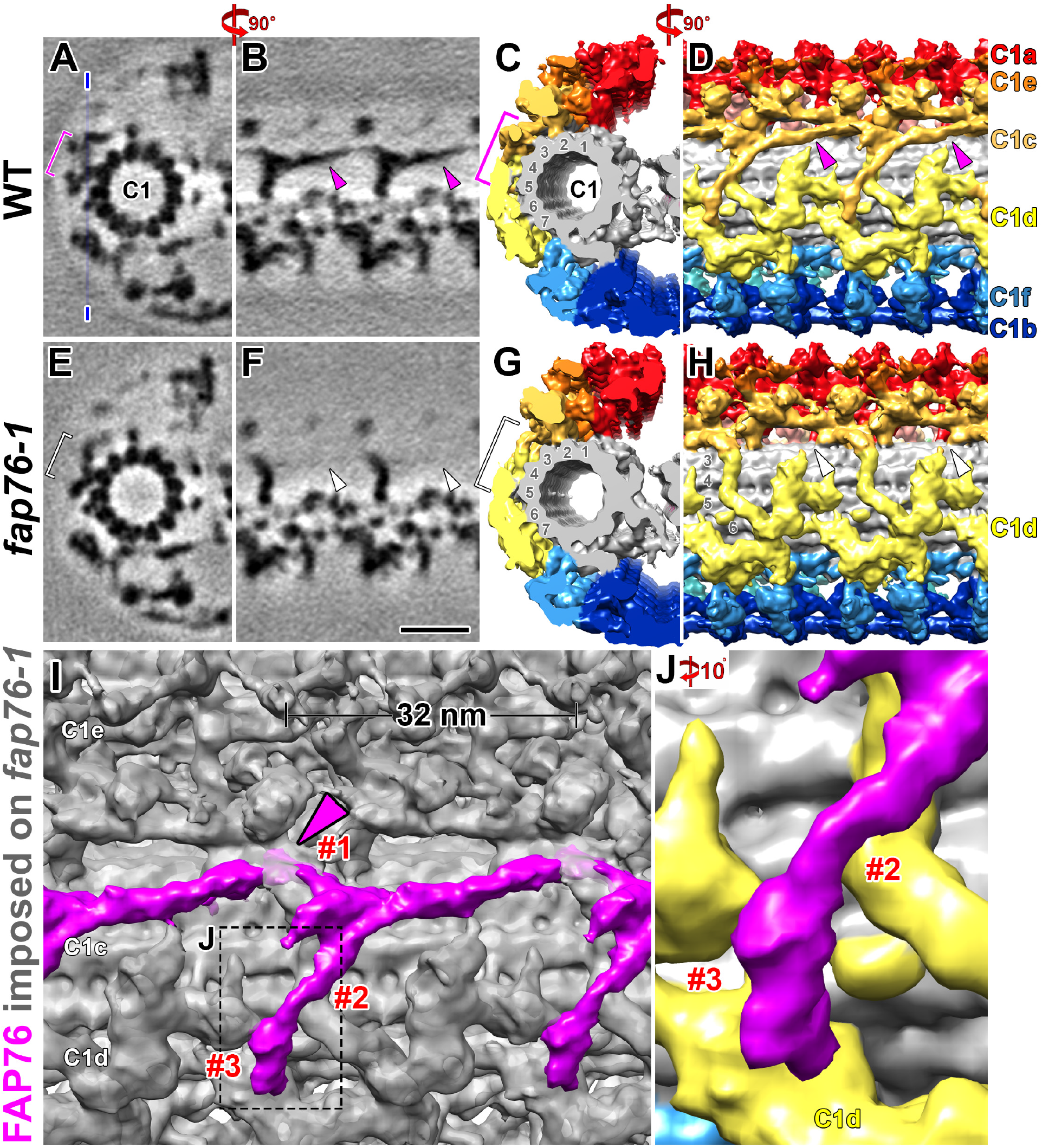
FAP76 is a C1c protein with multiple connections to neighboring structures. **(A-H)** The comparison of tomographic slices (A, B, E and F) and isosurface renderings (C, D, G, and H) between the averaged CA repeats of wild-type (A-D) and *fap76-1* axonemes (E-H) viewed in cross-sectional (A, C, E and G) and longitudinal (B, D, F and H) orientations, shows a triskelion-shaped structure that is present in the wild-type C1c projection (magenta brackets and arrowheads in A-D) but missing in *fap76-1* (white brackets and arrowheads in E-H). The thin blue line in A indicates the location of the tomographic slices shown in B and F. **(I-J)** The isosurface rendering of a difference map between the averaged CA of wild type and *fap76-1* shows the triskelion-shaped FAP76 densities (magenta) in longitudinal view superimposed on the averaged CA of *fap76-1* (grey) in overview (I) and zoomed-in (J, location indicated by box in I). Note the three connections (magenta arrowhead and #1-3) of FAP76 with neighboring structures, i.e. #1 with the C1c projection close to C1e (I), and #2 and #3 with C1d densities (yellow in J). Scale bar in F, 20 nm (valid for A, B, E and F).

The FAP76 density consists of a 32 nm long filamentous part and a thicker “branch” extending from the middle of the filament to connection #1 (Fig. 4 I). To analyze the FAP76 structure, we rescued *fap76-1* with N-terminally BCCP-tagged FAP76. Rescue cells swim (mostly) with wild-type motility (Fig. 3 A-C). Using immunofluorescence microscopy and SDS-PAGE, we confirmed that BCCP-FAP76 assembled into the axonemes of the rescued cells (Fig. 5 A and B). In the subtomogram averages of *fap76-1;BCCP::FAP76* axonemes labelled with streptavidin-nanogold, we found an additional density where the short “branch” of FAP76 connects to the C1e projection (Fig. 5 J-L, blue arrowheads), indicating the location of the FAP76 N-terminus. This additional density was not visible in wild-type axonemes or *fap76-1;BCCP::FAP76* without streptavidin-nanogold (Fig. 5 D-I, white arrowheads). Although the overall CA structure of *fap76-1;BCCP::FAP76* resembles that of wild type, one of the two globular-shaped structures at the C1c-e junction (C1c peripheral subunit 1 (psu1) and black arrow in Fig. 5 E and F) appears to be missing with and without incubation with streptavidin-Au (white arrow in Fig. 5 H, I, K and L). A classification analysis focused on C1c psu1 revealed that ∼40% of wild-type and ∼38% of *fap76-1* CA repeats, but less than 10% of CA repeats in the *fap76-1;BCCP::FAP76* rescued strain contained the C1c psu1 structure (Fig. 5). The close proximity of the FAP76 N-terminus to this subunit suggests that the BCCP tag in the rescue strain might disrupt assembly of the globular subunit. This defect might also explain the partial rescue of the photoshock response in *fap76-1;BCCP::FAP76* (Fig. 3 C). Comparisons of the proteomes of wild type, *fap76-1* (Table 1), and *fap76-1;BCCP::FAP76* (data not shown) did not reveal missing proteins that might correspond to C1c psu1, suggesting that C1c psu1’s apparent absence in the tomogram averages of the latter strain is due to positional flexibility that results when FAP76 is replaced with FAP76-BCCP.

**Figure 5.**
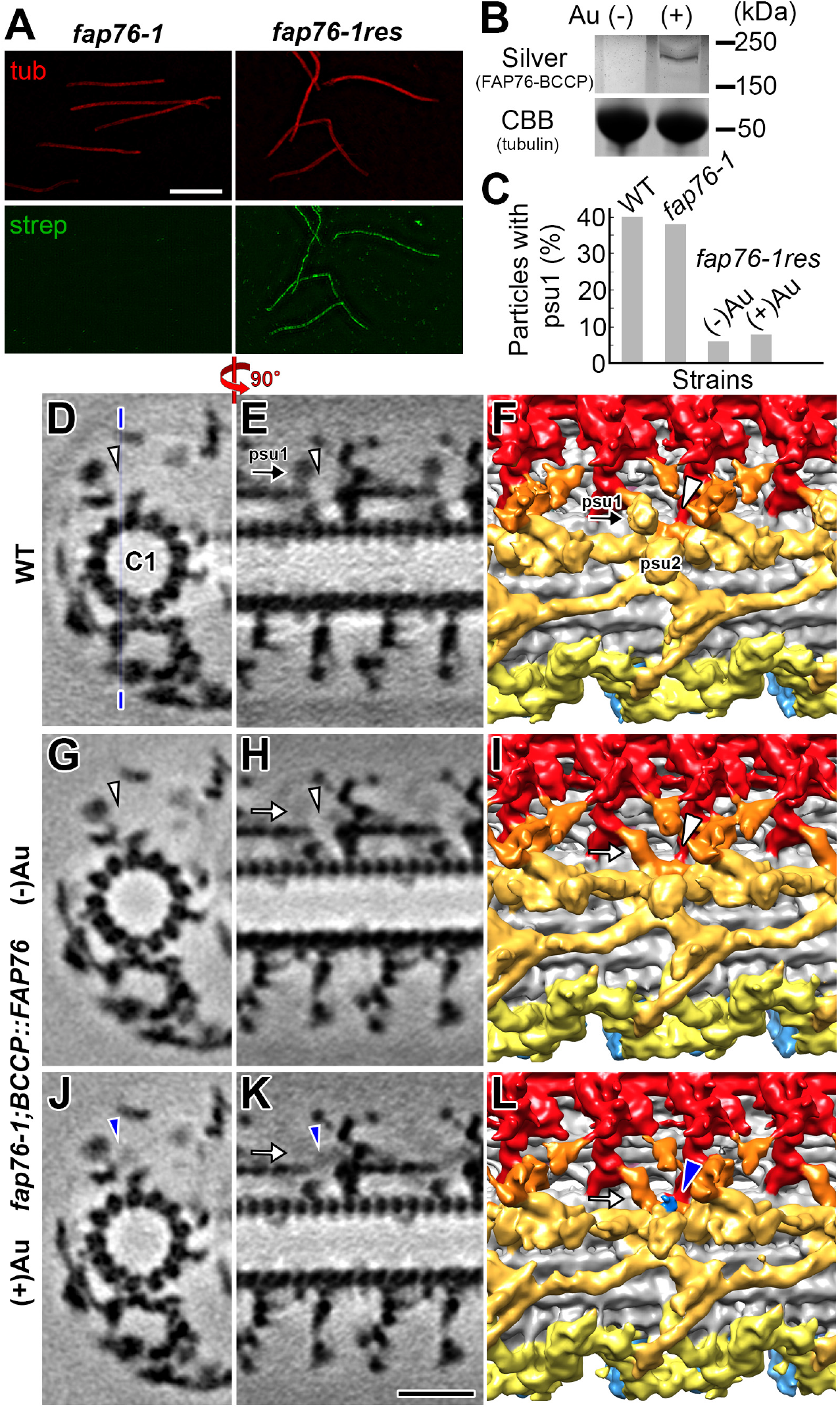
The FAP76 N-terminus is located at the interface between C1c and C1e projections. **(A)** Immunofluorescence light microscopy images of axonemes isolated from *fap76-1* (left) and *fap76-1;BCCP::FAP76* rescue (right) probed by anti-acetylated-tubulin antibody (red) and fluorescently tagged streptavidin (green). The streptavidin signal indicates that the BCCP-tagged FAP76 was assembled into the axoneme of the rescued strain. **(B)** SDS-polyacrylamide gel stained by a silver enhancement kit (top) to show that a specific band of appropriate relative mobility could be detected in the *fap76-1;BCCP::FAP76* axonemes treated with streptavidin-Au, but not in the control (-Au). Coomassie Brilliant Blue (CBB) staining (bottom) shows the tubulin bands as loading controls. **(C)** A classification analysis of the C1c peripheral subunit 1 (psu1/black arrows in E and F) in wild type and *fap76-1*, and *fap76-1;BCCP::FAP76* rescue axonemes (white arrows in H, I, K and L). The particle numbers (n) included in the averages for wild type, *fap76-1*, *fap76-1;BCCP::FAP76* (+Au) *and fap76-1;BCCP::FAP76* (-Au) were 664, 927, 1089 and 935 (see Table S3). **(D-L)** Comparison of tomographic slices (D, E, G, H, J and K) and isosurface renderings (F, I and L) of the averaged CA repeats of wild type (D-F) vs. *fap76-1;BCCP::FAP76* rescue strain either without (G-I) or treated with (J-L) streptavidin gold, viewed in cross-sectional (D, G and J) and longitudinal (E, F, H, I, K and L) orientations. The additional density of the streptavidin-gold label at the interface between the C1c and C1e projections in the gold-treated rescue strain (blue arrowheads in J-L) is not observed in wild-type or control CA (white arrowheads in D-I). Thin blue line in D indicates the location for the tomographic slices shown in E, H and K. Scale bar in A, 5 µm (valid for all fluorescence images); in K, 20 nm (valid for all EM images in D-L).

We also analyzed a second CLiP mutant, *fap76-2,* in which the *FAP76* gene was disrupted by insertion of the cassette at a location closer to the 3’end of the gene than in *fap76-1* (Fig. S4 A). The phenotype of *fap76-2* is very similar to that of *fap76-1* with regard to both motility (Fig. S4 B-D) and structural defects (Fig. S4 F and G). In contrast to *fap76-1*, MS analysis of *fap76-2* showed that a truncated FAP76 protein is assembled into the *fap76-2* axoneme, and that levels of two more proteins, DPY30 and FAP380, are reduced compared to *fap76-1* (Fig. S4 E-G). Competition with this truncated FAP76 protein could explain our observation that FAP76-HA expression in *fap76-2* results only in a partial rescue of motility (unpublished observations), whereas *fap76-1* is completely rescued (Zhao et al., 2019).

#### FAP81 is required for stable assembly of the C1e and C1c projections

*fap81* CA had the most severe structural defects of the four CLiP mutants. The wild type-mutant comparison revealed that the C1e and C1c projections were missing in the averaged *fap81* CA repeat (compare Fig. 6 A-C with D-F; Video 2), indicating that FAP81 is required for stable assembly of the C1e-c subcomplex. The estimated size of the wild-type C1e-c subcomplex is 1.2 MDa. However, MS and western blot analyses of *fap81* axonemes revealed that, of the 16 proteins associated with PF16, only FAP81 (172 kDa) and FAP216 (79 kDa) were substantially reduced (Table 1 and Fig S2 D). This suggests that some of the C1e-c proteins might still bind to the *fap81* axoneme but cannot be visualized in the averaged *fap81* CA due to partial reduction and/or positional flexibility. This is supported by the observation that in the region where FAP76 connects to the C1d projection (#3 connection, Fig. 4 I) a weak FAP76 density remained visible in the averaged *fap81* CA, whereas this density was missing in *fap76-1* and *fap76-1;fap81* (compare Fig. S5 A, C with B, D). A classification analysis focused on this region could clearly visualize the part of FAP76 that connects to the C1d projections in more than half of the *fap81* CA repeats (compare Fig. S5 E and F), which is consistent with MS data showing that a considerable number of FAP76 peptides were detected in the *fap81* axonemes (Table 1). Nonetheless, large parts of FAP76, including its N-terminal region, remained undetectable even in the class averages of *fap81*. This suggests that FAP81 interacts with the N-terminal part of FAP76, and without this interaction FAP76 becomes positionally flexible. Despite the partially remaining of FAP76 protein in *fap81*, the similar phenotypes of *fap81* and *fap76-1;fap81* indicate that the positionally flexible FAP76 did not function properly in *fap81* axonemes.

**Figure 6.**
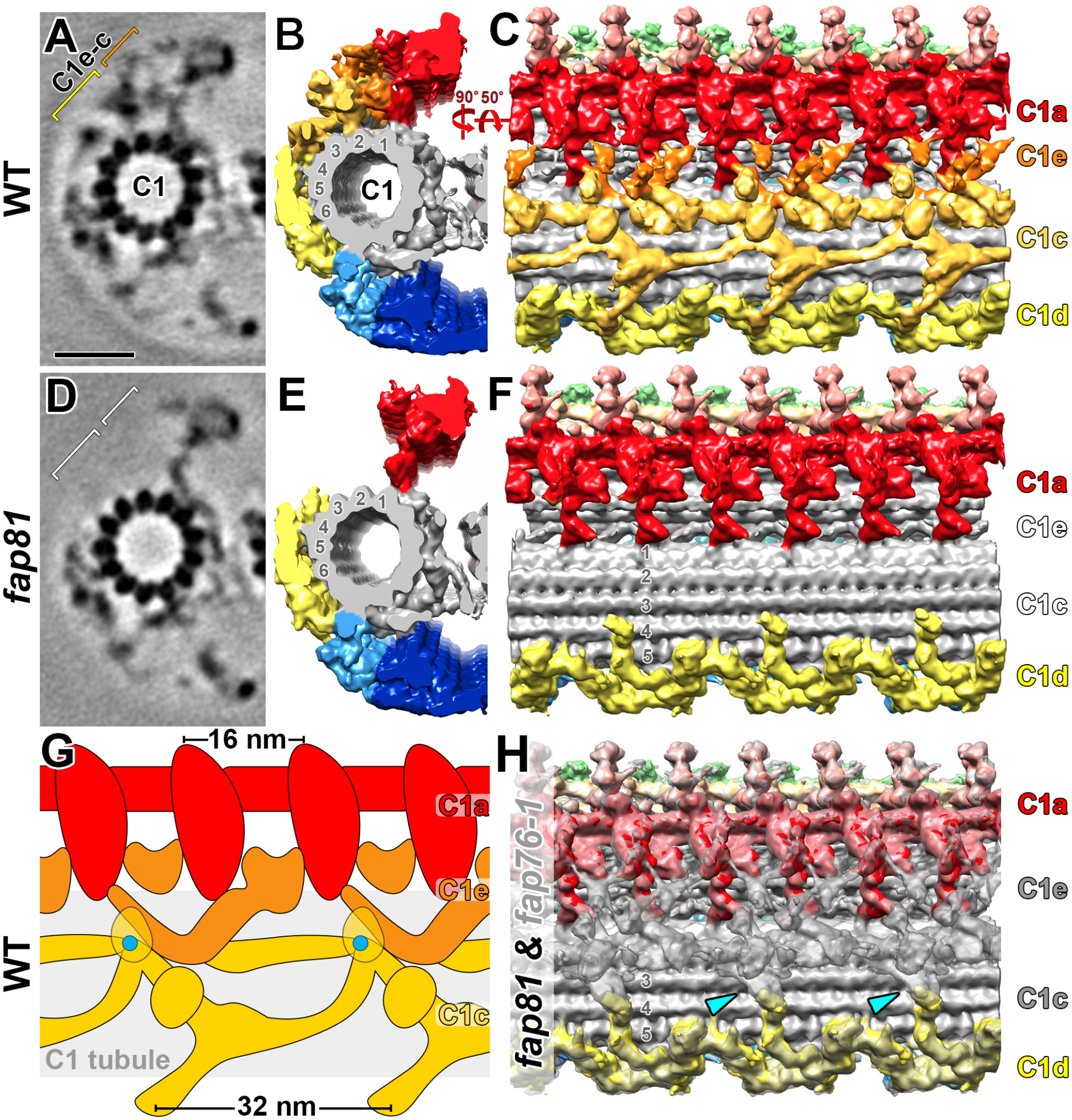
FAP81 is required for the stable assembly of the C1e-c subcomplex. **(A-F)** Comparison of tomographic slices (A and D) and isosurface renderings (B, C, E and F) between the averaged CA repeats of wild type (A-C) and *fap81* (D-F), viewed in cross-sectional (A and D) and longitudinal (B, C, E and F) orientations, shows that the C1e and C1c projections (orange/yellow brackets in A) are present in wild type but missing in *fap81* (white bracket in D). **(G)** Schematic drawing of the wild-type CA structure in longitudinal view to outline the densities of the C1a (red) and C1e (orange) projections and their interactions. Note the transition from 16 nm periodicity (C1a) to 32 nm periodicity (C1e). Blue dots indicate the locations of the FAP76 N-terminus. **(H)** An overlay of the averaged CA repeats of *fap76-1* (transparent gray) and *fap81* (colored) shows that the interaction (cyan arrowheads) between a rod-shaped C1d density and the gray C1c density, which remains present in the *fap76-1* CA and consists of at least the FAP81 protein. Scale bar: 20 nm (A valid for A and D).

Comparison of the wild-type and *fap81* CAs allowed a more precise characterization and delineation of the C1a and C1e densities. In wild type, the C1a projection, which repeats every 16 nm, has multiple interaction sites with the C1e projection, which repeats every 32 nm (Fig. 6 G). Our data clearly showed that alternating C1a projections interact with different parts of each C1c projection (Fig. 6 G). The results for *fap81* also provided insight into the interactions between FAP81 and C1d. Since FAP81 is not significantly reduced in *fap76-1* axonemes (Table 1), the remaining C1c densities in *fap76-1* are likely composed of FAP81 and FAP216. Comparison of *fap81* with *fap76-1* reveals that FAP81 interacts with a rod-shaped C1d density that is anchored to C1 protofilament 4 (Fig. 6 H).

#### FAP216, a C1c subunit, links the C1 microtubule and peripheral C1c projection densities

Our MS analyses revealed that, in addition to FAP81, FAP216 was significantly reduced in *fap81* axonemes, whereas FAP81 was present at wild-type level in *fap216* (Table I). This suggests that the assembly of FAP216 into the axoneme requires FAP81 but not *vice versa*. Structural analysis of *fap216* showed that two small densities were missing or reduced in the C1c and C1e projections, respectively (Fig. 7 E-H, light orange and white arrowheads). In the wild-type CA, the affected C1c structure links the more peripheral C1c projection density to the C1 microtubule between protofilaments 2 and 3 (Fig. 7 A-D, yellow arrowheads). The estimated molecular mass of the affected C1c density is ∼90 kDa, suggesting that this density is composed of a single copy of FAP216 (79 kDa). The affected C1e density, which forms a bridge between the C1c and the C1a projections in wild type (orange arrowhead in Fig. 7 A and C), is only partially reduced in *fap216* (light orange arrowhead in Fig. 7 E and G), and classification analysis revealed that this C1e density was missing in only 57% of the *fap216* CA repeats (Fig. S5 G-J). This, together with the results of the MS analyses, which did not detect any FAP216 peptides in *fap216* axonemes, indicates that the partial reduction of this density is likely a secondary defect. The protein(s) corresponding to the reduced C1e density could not be identified by our MS comparisons between wild-type and *fap216* mutant axonemes.

**Figure 7.**
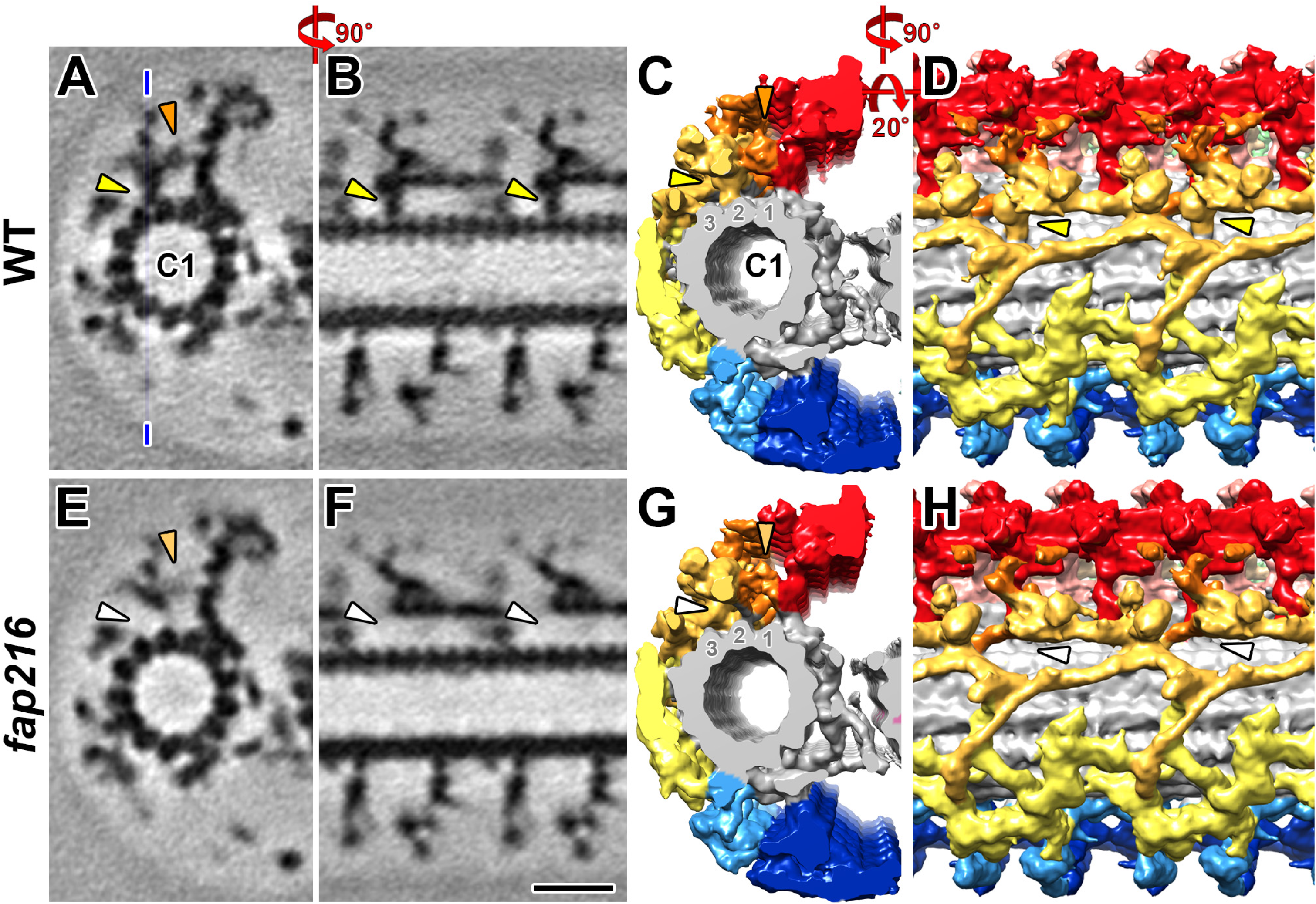
FAP216, a C1c protein that may bridge the C1 microtubule and peripheral C1c densities. **(A-H)** Comparisons of tomographic slices (A, B, E and F) and isosurface renderings (C, D, G, and H) between averaged CA repeats of wild type (A-D) and *fap216* (E-H) viewed in cross-sectional (A, C, E, and G) and longitudinal (B, D, F, and H) orientations shows that the *fap216* CA lacks a small C1c density (white arrowheads in E-H) that in wild type connects between C1 microtubule protofilament 2 and peripheral C1c densities (yellow arrowheads in A-D). In addition, part of the C1e projection (orange arrowheads in A and C) that bridges between the C1e-c subcomplex and the C1a projection is reduced in *fap216* (light orange arrowhead in E and F). Classification analysis confirmed that 57% of the *fap216* CA repeats lack this C1e density (see Fig. S4). Scale bar in F, 20 nm (valid for all EM images).

Although the structural CA defects in *fap216* are considerably less severe than those of *fap81*, both mutants display equally severe swimming defects. We also showed that transformation of *fap216* with the wild-type gene for FAP216 is sufficient to restore the wild-type phenotype (Fig. 3 A-C). Classification analyses of the *fap216* CA focused on the areas neighboring the structural defects did not reveal any positional flexibilities of the remaining densities, i.e. FAP216 does not appear to play a critical role in the stable assembly of the remaining C1a-e-c protein network. Instead, these results suggest that FAP216 is an essential component in a signal transduction cascade that regulates ciliary motility in *Chlamydomonas*.

## Discussion

### Composition and hierarchical assembly of the C1a-e-c supercomplex

Our wild type vs. mutant comparisons have revealed a 2 MDa PF16-associated protein network, the C1a-e-c supercomplex, within the ciliary CA. We show that PF16 is associated with at least 16 proteins (Table 1) with a combined molecular mass of 1.6 MDa, suggesting that multiple protein copies and/or additional proteins form the supercomplex. Our MS analyses also identified two proteins, FAP54 and FAP221, previously assigned to C1d, among the PF16-associated proteins, suggesting that they have stable attachments to C1a-e-c supercomplex proteins. Such strong interaction between the C1c and C1d projections is consistent with the observed structural connections.

We characterized mutant phenotypes and the molecular organization for five C1a-e-c supercomplex proteins. PF16 and FAP92 are C1a projection proteins, whereas FAP76, FAP81, and FAP216 are the first identified C1e and C1c proteins (Table 1 and Fig. 8; Video 2). The CA structural defects in the mutants suggest hierarchical assembly of the C1a-e-c supercomplex. The complex was disrupted in *pf16* axonemes suggesting that the C1a protein PF16 is a crucial scaffold protein required for stable assembly of the entire supercomplex (Figs. 2 and 8). Loss of PF16, and the supercomplex, destabilizes the entire C1 microtubule and associated projections, as only ∼10% of the *pf16* axonemes had “9+2” axonemes.

**Figure 8.**
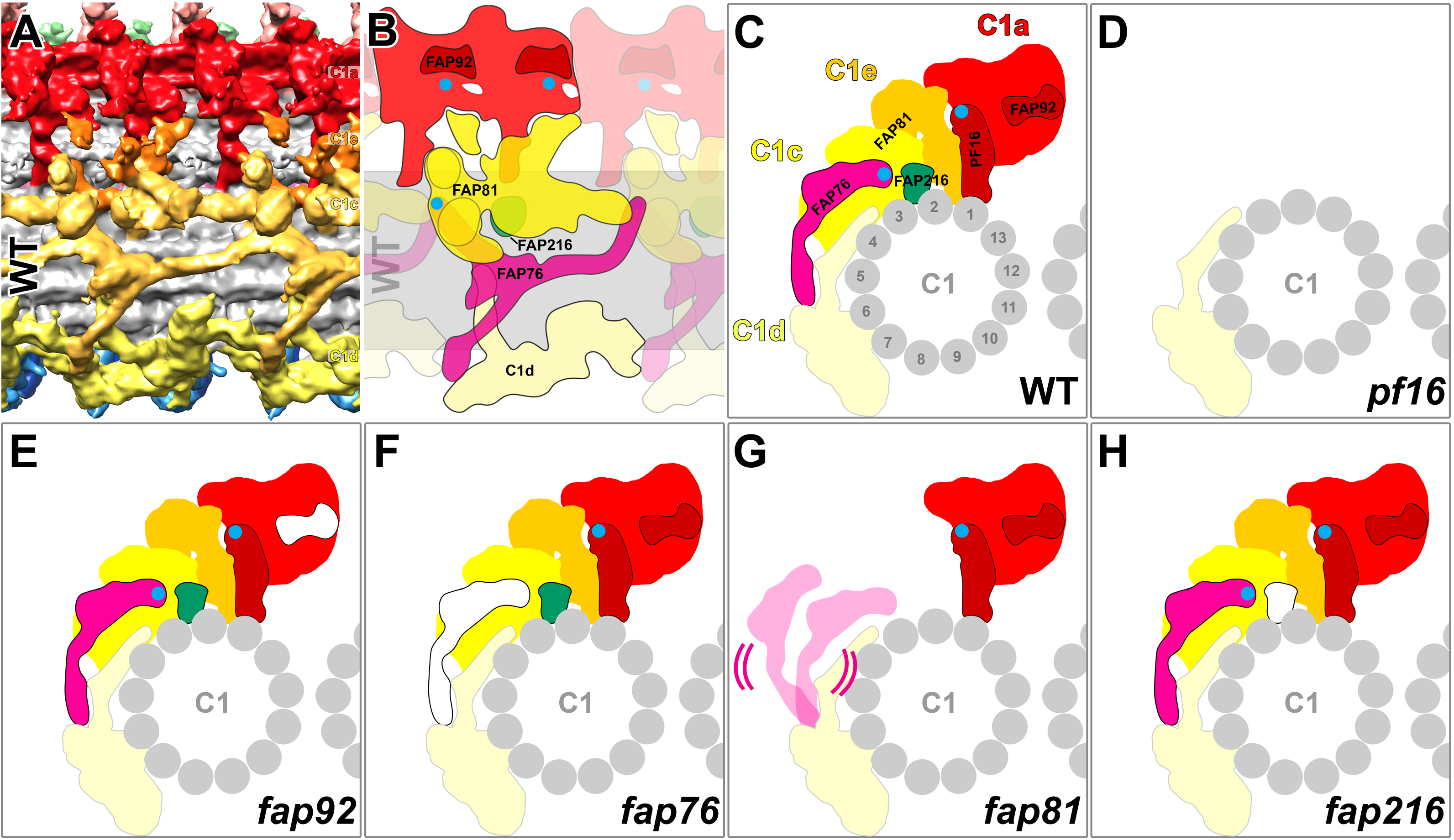
Models of the C1a-e-c supercomplex. **(A)** The isosurface rendering shows the 3D structure of the averaged *Chlamydomonas* wild-type CA repeat viewed in longitudinal orientation. **(B, C)** A schematic drawing of the 32 nm repeat of the *Chlamydomonas* wild-type CA in longitudinal orientation (B; same orientation as in A) and cross-sectional view (C) to highlight the protein compositions of the C1a-e-c supercomplex within the ciliary CA; this includes densities containing the subunits PF16 and FAP92 (dark red), FAP76 (magenta), FAP81 (yellow), and FAP216 (green). Blue dots indicate the C-terminus of PF16 and N-terminus of FAP76 proteins. **(D**-**H)** Schematic drawings of the 32 nm CA repeat of various C1a-e-c mutants in cross-sectional view to show the observed structural defects.

The second largest supercomplex defect was observed in *fap81* axonemes, suggesting that FAP81 is essential for stable assembly of the C1e-c subcomplex that consists of at least FAP76, FAP81 and FAP216. In contrast, the single loss of FAP76, FAP92 or FAP216 had minimal effects on the assembly of other C1a-e-c supercomplex components. Despite at least two FAP76 contact sites and one FAP81 contact site to C1d projection densities (Figs. 4 and 6), none of the mutants showed defects in C1d projection assembly. Likewise, classical thin-section TEM of two C1d-defective mutants, *fap46* and FAP74 RNAi, which lack the entire C1d projection, showed no obvious effects in structures corresponding to the C1a-e-c supercomplex (DiPetrillo and Smith, 2010; Brown et al., 2012). Our precise localization of FAP76, FAP81, FAP92, and FAP216 within the C1a-e-c supercomplex agrees well with the recent assignment for these proteins to the C1 microtubule (Zhao et al., 2019). However, previous immunoprecipitation experiments assigned FAP81 incorrectly to the C1a projection (Zhao et al., 2019), likely due to the tight association between the C1a and C1e-c projections (Fig. 6). This highlights that the integration of biochemical and structural data is critical for a comprehensive understanding of the molecular architecture of the CA.

### Supercomplex subunits form connections within the complex and with the C1d projection

The improved resolution of our cryo-ET data revealed that the C1a-e-c supercomplex makes three connections with the neighboring C1d projection and multiple interactions within the complex, such as the interface between the C1a and C1e projections, where the structural periodicity transitions from 16 nm (C1a) to 32 nm (C1e) (Fig. 6 G). Alternating C1a projections, which physically interact with each other, have different connections to each C1e projection, suggesting that the alternate C1a projections could have distinct function(s).

The C1a projection is anchored to C1 protofilament 1 with 16 nm periodicity (Figs. 2 C, 6 E), which might involve the N-terminal domain of PF16 (Fig. 8). Every 16 nm, the C1e-c subcomplex has two adjacent connections to C1 protofilament 2. The smaller connecting density consists of FAP216 (yellow arrowheads in Fig. 7 A-D) and did not appear essential for complex docking, whereas the larger distal connection is a protein interaction hub involving multiple structures, including C1e, the C1c peripheral subunits 1 and 2, FAP81, and the N-terminal domain of FAP76 (Fig. 4 I, #1 connection). In addition to being part of the latter interaction hub, the triskelion-shaped FAP76 formed at least three more interfaces with neighboring structures: Connections to each other in the longitudinal direction (Fig. 4 I), and two connections to the neighboring C1d projections (Fig. 4 J). The C1d projections themselves were bound every 32 nm to C1 protofilaments 4 and 6 (Figs. 4 G and H; 6 E and F; 8). Given the motility defects observed in all C1a-e-c mutants, including *fap216* with its very small structural defect, these subunits are likely involved in the same signal transduction pathway that regulates ciliary motility. Their connectivity with neighboring structures may provide the structural basis for signal transmission.

### The C1e-c subcomplex, likely with C1d, forms a signaling pathway regulating ciliary motility

Most previously identified CA protein mutations, including the C1a mutant *pf16*, result in paralyzed or twitching-only cilia (Dutcher, 1984; Smith and Lefebvre 1996; Rupp et al., 2001). The four C1a-e-c supercomplex mutants identified here, *fap76-1*, *fap81, fap92,* and *fap216*, have slow swimming phenotypes. However, loss of FAP92 did not result in major motility or structural defects, likely because the major C1a components remained largely unaffected. In contrast, cells of the remaining three mutants, *fap76-1*, *fap81,* and *fap216*, frequently changed swim orientations, resulting in curving swimming paths (Fig. 3 B). A straight swimming orientation, or sharp turns, seen in *Chlamydomonas* cells, are coupled to synchronous or asynchronous ciliary beating, respectively (Polin et al., 2009). Consistent with this, the period of synchronous beating between the two flagella of the mutant cells was shorter than that of wild type (Video 3), causing frequent reorientation during swimming. If the overall shape of the flagellar waveform is not changed, as is the case here (Video 3), then the loss of synchrony between the flagella could be caused by random delays in switching of dynein activity from one side of the axoneme to the other in one of the two flagella. Our findings indicate that the C1e-c subcomplex, which is largely composed of FAP76, FAP81 and FAP216, may play a critical role in coordinating the oscillatory switching of dynein activity, and thus in maintaining synchrony between the two beating cilia.

The C1e-c subcomplex may act in the same regulatory pathway as the adjacent C1d projection, because previous studies reported similar motility defects for *Chlamydomonas* C1d mutants, i.e. slow swimming, uncoordinated ciliary beating, and a deficient photoshock response (DiPetrillo and Smith, 2010; 2011; Brown et al., 2012). In our model the C1e-c subcomplex is part of the protein network that requires PF16 for stable assembly, but it has a 32 nm periodicity, identical to that of the C1d projection, and twice that of the C1a projection. Our cryo-ET data revealed that the C1c projection interacts directly with the C1d projection through FAP76 and FAP81 (Fig. 8 B), and our combined sucrose gradient and MS data provided evidence that the PF16-associated protein network is physically attached to at least two proteins previously assigned to C1d. Thus, the C1e-c subcomplex may be physically and functionally distinct from the C1a projection, but closely related to C1d. Given the more severe motility defects observed in the C1d mutants (DiPetrillo and Smith, 2010; 2011; Brown et al., 2012), regulatory signals might be transmitted from the C1d projection to the C1e-c subcomplex through their direct connections. From the C1e-c subcomplex the signal could be sent downstream either to the C1a projection or to radial spoke heads to ultimately modulate dynein activities. One such signal could be initiated by changes in intraflagellar calcium concentration, because the C1d protein FAP221 binds calmodulin in a Ca^2+^-sensitive manner, and thus could mediate Ca^2+^-induced changes in flagellar waveform (Witman, 1993; DiPetrillo and Smith, 2010; Brown et al., 2012). Although FAP76 could still bind to the C1 microtubule and connect to the neighboring C1d projection in the *fap81* mutant (Fig. S5 A-F), the positional flexibility of FAP76 caused by loss of its N-terminal binding partner FAP81 could be sufficient to disrupt the proposed signal-transmission cascade between C1d and C1c-e.

### Spatial association between the CA and the DMT-associated dynein-regulators

How the CA signals are transmitted to DMT dyneins through the interaction with RS is poorly understood. In *Chlamydomonas*, the CA is slightly twisted around it’s longitudinal axis (90° twist over 2 µm CA length; Carbajal-González et al., 2013) and the CA has been shown to rotate around the ciliary axis during motility (Kamiya, 1982). Therefore, the CA projections in *Chlamydomonas* flagella are not expected to have a fixed interaction with radial spokes from a specific DMT. However, previous *in vitro* microtubule sliding assays and classic thin-section EM showed that the C1 microtubule was predominantly oriented toward the set of actively sliding DMTs, which switches between DMTs 2-4 and DMTs 6-8 in the two halves of the axoneme (Wargo and Smith, 2003). In addition, the orientation of the C1 microtubule seems to be correlated with the flagellar bending direction as observed in axonemes prepared by freeze-etching (Mitchell, 2003). Switching of ciliary bending direction, which is critical to generate the undulating motion typical for cilia, correlates with the switching of dynein activity that drives inter-doublet sliding between either DMTs 2-4 or DMTs 6-8 (Wais-Steider and Satir, 1979; Tamm, 1984; Sale, 1986; Hayashi and Shingyoji, 2008; Lin and Nicastro, 2018). Therefore, we examined the spatial association between the CA projections and DMTs in wild-type and mutant axonemes. We found that the C1a-e-c supercomplex predominantly faced DMTs 6-8 in isolated and inactive wild-type and C1e-c mutant axonemes (Fig. S3 H). The preferential orientation of the C1a-e-c supercomplex towards DMTs 6-8 is consistent with the CA projections playing a role in controlling ciliary bending direction. However, the axonemes we isolated were inactive (no ATP added in the buffer), therefore, all dyneins were in their post-power stroke conformation. Future studies of the CA in actively beating cilia could reveal if the relative location of the C1a-e-c supercomplex switches between the DMTs 2-4 or DMTs 6-8 in a bend-direction specific manner.

### FAP216 plays a role in chemical signal transduction between CA and radial spokes

A previous study reported that the motility defects of the *pf6* mutant, which lacks most of the C1a-e-c projections (data not shown), could be restored by adding exogenous tags to the radial spoke heads (Oda et al., 2014). This result led to a working model that the CP–RS communication is, at least in part, mediated by nonspecific collision-based mechano-signaling between the CA projections and radial spoke heads without requiring specific physiological protein-protein interactions at the CP–RS interface (Oda et al., 2014). The severe motility defects of the *fap216* mutant suggest that a mechanically intact CA–RS interface alone is not sufficient for proper CA–RS communication and regulation of dynein activity. Despite only missing a small density in the center of the CA, 18 nm away from the interface with the radial spoke heads (Figs. 7 G and 8), the motility defect in *fap216* is as severe as in *fap81*, which lacks the entire C1e-c subcomplex. This raises the possibility that FAP216 contributes to a chemical signal transduction pathway between the CA and radial spokes to regulate ciliary motility.

In humans, many axonemal gene mutations of have been shown to cause ciliopathies, such as PCD (Horani and Ferkol, 2018). Historically, conventional TEM analysis of patient ciliary samples is considered a standard clinical diagnostic tool for PCD (Kott et al., 2013; Khouri et al., 2016; Edelbusch et al., 2017). However, the resolution of conventional TEM is greatly limited and could not visualize the defects in the *fap216* mutant, hindering the diagnosis of PCD-types caused by minor structural alterations. Therefore, future comparisons of normal and potential ciliopathy CA structures by cryo-ET may facilitate recognition of previously undiagnosed or misinterpreted defects in human patients (Lin et al., 2014). Overall, our study provides a partial “molecular blueprint” for the C1a-e-c supercomplex that will be the foundation for future studies into detailed protein-protein interactions and molecular mechanisms by which CA signals are transmitted to the radial spoke heads to ultimately regulate dynein activity and thus ciliary beating.

## Materials and methods

### Strains and culture conditions

*Chlamydomonas* wild-type strains used were g1 (*nit1*, *agg1*, *mt+*, a cross of *nit1-305* to CC-124, Pazour et al., 1995; *Chlamydomonas* Resource Center, https://www.chlamycollection.org<u>, CC-5415</u>) and A54-e18 (<u>CC-2929</u>). The mutant *pf28* (CC-3661) was obtained from *Chlamydomonas* Resource Center and *pf16* (D2, the *pf16* insertional allele) was generated as previously described (Smith and Lefebvre, 1996). The *pf28* and *pf16* (D2) strains were mated to generate *pf28;pf16* double-mutant (DiBella et al., 2004). Strains from the *Chlamydomonas* Library Project (CLiP; https://www.chlamylibrary.org; Li et al., 2016) included the parental strain CC-5325 and the insertional mutants *fap76-1* (CLiP ID: LMJ.RY0402.089534), *fap76-2* (LMJ.RY0402.088713), *fap81* (LMJ.RY0402.092632), *fap92* (LMJ.RY0402.204383) and *fap216* (LMJ.RY0402.218389), all of which were obtained from the *Chlamydomonas* Resource Center. For MS analysis of axonemal proteins, *fap76-1* was crossed to g1 (Zhao et al., 2019), which served as the control. As previously described (Fu et al., 2018), *Chlamydomonas* cells were maintained in solid Tris-acetate-phosphate (TAP) plates (supplied with 7.5 µg/ml paromomycin for CLiP mutants) and cultured in liquid TAP medium or modified M medium (Witman, 1986) under a 12:12 h light:dark cycle at 23°C with filtered air bubbling into the growing culture. Insertion sites of CLiP mutants were confirmed by PCR (Fig. S2 A) using the primers indicated in Table S4.

### Generation of tagged strains and rescue of mutants

To generate the BCCP-tagged *pf16;PF16::BCCP* strain, the BCCP domain of *Chlamydomonas* acetyl-coenzyme A carboxylase gene (Cre17.g715250, Phytozome 12; Amino acids 141-228 were used for tagging) was amplified by RT-PCR from genomic DNA from wild-type *Chlamydomonas* (strain A54-e18, CC-2929) as previously described (Oda and Kikkawa, 2013). A BCCP-tagged *PF16* construct was generated by ligating the BCCP domain into the C-terminal MluI site of the pB6D2 plasmid that contains the *PF16* gene (a 4.5 kb genomic fragment in pBluescript that rescues the mutant phenotype, Smith and Lefebvre, 1996). The BCCP-pB6D2 plasmid was co-transformed into *pf16A* or D2 (a *pf16* insertional allele), along with the *APHVIII* gene, using the glass bead method (Kindle 1990). Transformed cells were plated onto TAP plates with 20 µg/ml paromomycin. Cells were screened first for rescued motility, then by western blot of axonemes to confirm the presence of PF16 along with the BCCP tag (rabbit anti-PF16 antibodies; 1:1000 affinity purified; Smith and Lefebvre, 1996) and a streptavidin-HRP probe (1:5000, GE Healthcare No. GERPN1231).

For the FAP76-BCCP rescue construct, sequence encoding the BCCP tag was amplified from pIC2L-BCCPC-3xHA-Hyg (kindly provided by Professor Toshiyuki Oda, University of Yamanashi, Japan) with primers F7/R7 and then was inserted into the MauB1 site of the plasmid pBC8, which contains a hygromycin cassette and the complete *FAP76* gene (Cre09.g387689, Phytozome 12), to yield pBC25.

For the FAP216-BCCP rescue construct, portions of the *FAP216* gene (Cre12.g497200, Phytozome 12) were amplified from genomic DNA with primers F8/R8, F9/R9, F10/R10. Plasmid pBH (Zhao et al., 2019) was linearized by digestion with NdeI and SbfI, and the *FAP216* fragments then assembled into it using NEBuilder HiFi DNA assembly master mix (NEB), yielding pBC26. To introduce the BCCP tag, sequence encoding the tag was amplified from pIC2L-BCCPC-3xHA-Hyg with primers F11/R11 and then was cloned into pBC26 at the MauB1 site, yielding pBC27.

All constructs were verified by sequencing. *Chlamydomonas* cells were transformed by the glass-bead method (Kindle, 1990). After transformation, cells were grown on TAP agar supplemented with 10 µg/ml hygromycin (Sigma-Aldrich). Cells were screened for rescued motility; incorporation of the construct was then confirmed by western blotting with a streptavidin-HRP probe (1:3000, Molecular Probes). West Dura Extended Duration Substrate (Thermo Fisher Scientific) was used for HRP detection.

### RNA isolation and RT-PCR

Total RNA was isolated from wild-type and *fap92* cells using RNeasy Plus Mini Kit (Qiagen). cDNA synthesis was performed using SuperScrip IV Reverse Transcriptase (Invitrogen) following the manufacturer’s protocol, and both Oligo(dT)_20_ and random hexamers were used as primers. *FAP92* (Cre13.g562250, Phytozome 12) fragments were then amplified using primers F12/R12 (Table S4) and sequenced. The gene encoding G protein β subunit (Schloss, 1990) was used as the control and amplified with primers F13/R13 (Table S4).

### Sucrose gradient analysis

Isolated axonemes were extracted in 0.5 M KI in NaLow (10 mM HEPES pH 7.5, 5 mM MgSO_4_, 1 mM DTT, 0.5 mM EDTA and 30 mM NaCl) at 6 mg/ml for 30 min on ice. After centrifugation, the extracted proteins were dialyzed into NaLow and then were loaded onto a 5-20% sucrose gradient made in NaLow buffer and centrifuged at 35,000 rpm for 16 h in a SW41Ti rotor (Beckman Coulter). Gradients were fractionated (0.5 ml fractions) from bottom (fraction 1) to top (fraction 24).

For western blot analyses, the sucrose gradient fractions were separated using SDS-PAGE and transferred to polyvinylidene difluoride (PVDF) membrane. For detection of PF16 protein, membranes were probed with anti-PF16 antibodies (1:1000 affinity purified). The ECL Prime Kit (GE Healthcare) was used for HRP detection.

### Immunofluorescence microscopy of *fap76-1* and *fap76-1;BCCP::FAP76*

Immunofluorescence microscopy was performed as described previously (Zhao et al. 2019). Briefly, the slides with attached intact axonemes were treated overnight at 4°C with blocking buffer containing the diluted primary antibody (mouse anti-acetylated tubulin, Sigma-Aldrich, clone 6-11B-1, 1:1000). On the next day, the slides were washed 4 times over 1 h with blocking buffer, and then treated for 1 h with blocking buffer containing the secondary antibody (F(ab’)2-goat anti-mouse IgG (H+L) cross-adsorbed secondary antibody, Alexa Fluor 568, Invitrogen, A11019, 1:1000) and Streptavidin (Alexa Fluor 488, Invitrogen, A32354, 1:200). The samples were mounted and examined as before (Zhao et al., 2019) using structured illumination microscopy (SIM) carried out on a DeltaVision OMX system (GE Healthcare) with a 1.42 NA 60x Plan-Apochromat objective lens (Olympus) and immersion oil with a refraction index of 1.512. SIM images were reconstructed with softWoRx 6.1.3 (GE Healthcare). Capture times and adjustments were the same for images with the same antibody. Image brightness and contrast were adjusted using ImageJ (National Institutes of Health).

### Analysis of motility phenotypes

All observations and recordings were carried out on cells grown in liquid TAP medium at room temperature. The swimming speed and swimming paths of cells were analyzed as described previously (Zhao et al., 2019). To analyze swimming speed, 50 µl of cell culture were transferred to a plastic chamber (0.127-mm deep Fisherbrand UriSystem DeciSlide; Thermo Fisher Scientific, Waltham, MA). Cells were imaged with non-actinic (deep-red) light using a Zeiss inverted microscope equipped with a 16x/0.35 NA Plan objective and a Kopp #2408 red long-pass filter (Kopp Glass Inc., Pittsburgh, PA). Movies were recorded at 30 images/s with a digital charge-coupled device (CCD) camera (UP-610; UNIQ Vision, Santa Clara, CA) and Video Savant 3.0 software (IO Industries, London, ON, Canada). Swimming speeds were determined using Image J software as previously described (Awata et al., 2015). To record swimming paths, 1-s exposures were acquired using white light and phase-contrast optics on a Zeiss Axioskop 2 Plus microscope equipped with a 20x Plan-NEOFLUAR 0.5 NA Ph2 objective, a digital CCD camera (AxioCam MRm), and AxioVision 3.1 software (Zeiss).

For video recording of swimming cells, a pco. 1200 hs camera with Camware software (the Cooke Corporation, Londonderry, NH) was used (500 frames per second) on an Axioskop microscope (Zeiss), and videos were taken with a 40x/0.65 NA objective lens under phase-contrast mode. Cells were in TAP medium at room temperature. A red filter placed on the light source was removed to induce waveform switching due to photoshock. Videos (saved at 20 fps) were created in ImageJ (Schneider et al., 2012) and the videos of different swimming cells were combined in Photoshop to generate Video 3. To evaluate photoshock ability at least 30 cells were analyzed.

### Axoneme preparation

Axonemes of *Chlamydomonas* cells were purified by the pH shock method as previously described (Witman, 1986; Song et al., 2015). Briefly, cells were cultured in liquid TAP medium, collected by centrifugation (2200 rpm for 5 min), and washed twice with fresh M-N/5 minimal medium (Iomini et al., 2009). The cell pellet was resuspended in pH shock buffer (10 mM HEPES, 1 mM SrCl_2_, 4% sucrose, 1 mM DTT, pH 7.4), then 0.5 M acetic acid was added to the buffer to reduce the pH to 4.3. After 80 sec, 1 M KOH was added to increase the pH to 7.2; the pH shock treatment was performed on ice. After pH shock, 5 mM MgSO4, 1 mM EGTA, 0.1 mM EDTA and 100 µl protease inhibitor (Sigma-Aldrich) were added to the solution. The solution was centrifuged (1800 g for 10 min, 4°C) to separate the detached flagella from the cell bodies. To further purify the flagella, the flagella-containing supernatant was centrifuged twice with a 20% sucrose cushion (2400 g for 10 min, 4°C). After centrifugation, 1% IGEPAL CA-630 (Sigma-Aldrich) was added to the supernatant to demembranate the flagella for 20 min at 4°C. Axonemes were collected by centrifugation (10,000 x g for 10 min, 4°C), and the freshly isolated axonemes then resuspended in HMEEK buffer (30 mM HEPES, 25 mM KCl, 5 mM MgSO_4_, 0.1 mM EDTA and 1 mM EGTA, pH 7.2). Axonemal samples were either plunge-frozen for cryo-ET or stored at −80°C for biochemical assays.

### Streptavidin gold labeling

For gold-labeling of axonemes from *pf16;PF16::BCCP* and *fap76-1;BCCP::FAP76*, 5 µl of 80 µg/ml 1.4-nm-sized streptavidin nanogold particles (Nanoprobes, Inc.) was added to 200 µl of the freshly prepared axonemal solution and incubated for 4 h at 4°C. Meanwhile, 200 µl of samples without added streptavidin gold particles were also prepared as control. The axonemes were washed by adding 800 µl HMEEK buffer and then collected by centrifugation (10,000 g for 1 min, 4°C) and resuspended in HMEKK buffer. Axonemal samples treated with and without streptavidin gold were plunge-frozen for cryo-ET analysis. In addition, 40 µg of the axonemes from the *fap76-1;BCCP::FAP76* sample (with and without gold particles) were separated on a 4-12% gradient SDS-polyacrylamide gel. The gel was stained with a silver enhancement kit (Nanoprobes, Inc.) for 45 min at room temperature and the bands then imaged using a ChemiDoc Touch Imaging System (Bio-Rad). The gel was then stained with Coomassie brilliant blue for 2 h and de-stained until the background was clear.

### LC-MS/MS

Axonemal protein (40 µg) of the wild type, *fap76-2*, *fap81* and *fap216* strains were separated on a 4-12% gradient SDS-polyacrylamide gel. After the proteins had entered the gel 3.0 - 3.5 cm, the gel was stained with Coomassie brilliant blue for 30 min and de-stained until the background was clear. Each gel lane was cut into 4 slices and each slice was further excised into 1-mm cubic pieces. In-gel trypsin digestion and peptide identification were conducted by the proteomics core facility at UT Southwestern Medical Center. Proteins that had more than 10 unique peptides identified in the wild-type sample were selected for further analyses. Quantification of the mutant:wild-type ratio for each axonemal protein was estimated by the MIC Sin value (Trudgian et al., 2011). Data for *pf16* are from Zhao et al., (2019; Repeat 2, Table S4). Some data for *fap76-1* are from Zhao et al. (2019; Repeat 2, Table S6). Intensity-based absolute quantification (IBAQ) (Schwanhausser et al., 2011) and Top3 precursor quantification methods (Top3) (Silva et al. 2006) were used to estimate the abundance of each protein.

For MS studies of sucrose gradient fractions from *pf16* (D2) or *pf28;pf16*(D2) axonemal extracts, proteins were run 0.5 cm into a 10% polyacrylamide resolving gel and stained with Coomassie blue. The protein-containing segments were excised and analyzed by matrix-assisted laser desorption/ionization time-of-flight (MALDI-TOF) tandem MS with post-source decay at the University of Massachusetts Medical School.

### Cryo-ET

Freshly prepared axonemal samples (30 µl) were gently mixed with 10 µl of 10-fold-concentrated, BSA-coated 10 nm gold solution (Iancu et al., 2006). 4 µl of the solution was applied to a glow-discharged (30 s at −35 mA) copper R2/2 holey carbon grid (Quantifoil Micro Tools GmbH, Jena, Germany). After removing excess liquid by blotting the grid from the back side with Whatman filter paper No. 1 for 2 s, the grid was plunge-frozen into liquid ethane using a homemade plunge-freezer. Grids were then stored in liquid nitrogen until used.

For most strains, grids were mounted in Autogrids^TM^ (Thermo Fisher Scientific, Hillsboro, OR) and loaded into a Titan Krios transmission electron microscope (Thermo Fisher Scientific, Hillsboro, OR) operated at 300 kV. The microscope control software SerialEM (Mastronarde, 2005) was used to acquire tilt series images in low-dose mode from −60° to 60° with 1.5-2.5° increments using a dose-symmetric tilting scheme (Hagen et al., 2016). The images were recorded with a 4k X 4k K2 direct electron detection camera (Gatan, Pleasanton, CA) in counting mode (15 frames, 0.4 s exposure time per frame and a dose rate of 8 electrons/pixel/s for each tilt image). The post-column energy filter (Gatan) was operated in zero-loss mode (20-eV slit width) and a Volta-Phase-Plate (Danev et al., 2014) were used with −0.5 µm defocus. The magnification was set to 26,000 with an effective pixel size of 5.5 Å. The total electron dose per tilt series was limited to ∼100 e/ Å^2^.

Grids of the *pf16*, *pf16;PF16::BCCP* and *pf6;pf16;PF16::BCCP* samples were imaged using a Tecnai F30 transmission electron microscope (Thermo Fisher Scientific, Hillsboro, OR) operated at 300 kV. A 2k X 2k charge-coupled device camera (Gatan, Pleasanton, CA) was used for recording tilt series images with SerialEM (Mastronarde, 2005) as described above, except neither dose-symmetric tilting nor Volta-Phase-Plate were used, and the defocus was set to −8 µm. The magnification was 13,500 with an effective pixel size of 1.077 nm.

### Image processing

Frame alignment for data collected on the K2 camera, alignment of tilt serial images and tomogram reconstruction were performed as previously described (Fu et al., 2018). In brief, motion correction of the frames was done with a script extracted from the IMOD software (Kremer et al., 1996). The IMOD software was also used for aligning the tilt serial images using the 10 nm gold particles as fiducial markers and for reconstructing the tomograms by weighted back-projection. For subtomogram averaging either CA or DMT repeats were picked from the raw tomograms, and the alignment and missing-wedge compensated averaging was performed using the PEET software (Nicastro et al., 2006), which is integrated in the IMOD software package. The axoneme and CA orientation (proximal to distal) was determined for each tomogram based both the DMT orientation in the axoneme cross-section (e.g. “clockwise” orientation as shown in Fig. 1H represents “viewed from distal”) and on initial averages that included only the CA repeats within each tomogram. After the CA polarity and the same center of the repeat was determined for each tomogram, a second alignment was performed combining the CA repeats from all tomograms in the correct orientation and periodicity register. After global alignment of all CA repeats, the alignment of each individual CA microtubule was refined by local alignment of each microtubule and its associated projections separately while the other microtubule was masked as described previously (Carbajal-González et al., 2013). Visualization of the three-dimensional structures of the averaged CA and DMT repeats was done with the UCSF Chimera package software (Pettersen et al., 2004). For generating the isosurface rendering images the same isosurface threshold was applied to the wild type and mutant averages. Classification analyses used a principal component analysis clustering method incorporated in the PEET software (Heumann et al., 2011). The number of tomograms and CA repeats, as well as the estimated resolution of the averages (using the 0.5 criterion of the Fourier shell correlation) are summarized for each strain in Table S3.

### Online supplemental material

Fig. S1 shows different conformational states of the C2a projection and refined structural details within the C1a-e-c supercomplex. Fig. S2 shows the genotyping analysis by PCR of the *fap76-1*, *fap76-2*, *fap81*, *fap92,* and *fap216* CLiP mutants to confirm disruption of the corresponding genes; immunoblot analysis of PF16 in *pf16* rescue strain and seven of the strains studied. Fig. S3 shows the isosurface renderings of averaged 96 nm axonemal DMT repeats from wild type and the CLiP mutants, confirming that the mutants have no DMT-associated defects; the orientation of the C1a-e-c supercomplex relative to specific DMTs is also shown. Fig. S4 depicts insertion sites for the *fap76* mutant alleles and shows the phenotype analyses and structural defects of the *fap76-2* mutant. Fig. S5 provides classification analyses of the *fap81* and *fap216* mutant showing that half of the *fap81* CA particles contain FAP76 density, and that the C1e density was reduced in the *fap216* mutant.

Table S1 lists all the proteins that co-sedimented with PF16 in our sucrose gradient analysis and that were significantly reduced in *pf28;pf16* axonemes as revealed by sucrose gradient and MS analyses. Table S2 provides unique peptide numbers and mutant/wild-type ratios of the C1 proteins (Zhao et al., 2019) in the C1a-e-c mutants studied here. Table S3 summarizes the image processing information (# of tomograms, CA particles, and resolution) for the strains used in this study. Table S4 lists the primers used in this study.

Video 1 shows the three-dimensional isosurface rendering of the averaged *Chlamydomonas* wild-type CA repeat. Video 2 summarizes the location and structural organization for FAP76, FAP81, FAP92 and FAP216 within the ciliary CA structure. Video 3 is a combination of videos showing swimming cells of *Chlamydomonas* wild type and the mutant strains studied here.

## Acknowledgements

We thank T. Loreng of Dartmouth College for technical assistance, and K. Cai and L. Gui of UT Southwestern Medical Center (UTSW) for discussions and critical reading of the manuscript. We thank A. Lemoff and the proteomics core facility at UTSW and M. Dubuke of the MS facility at the University of Massachusetts Medical School for the LC-MS/MS analyses. We are grateful to D. Stoddard for management of the UTSW cryo-electron microscope facility, which is funded in part by a CPRIT Core Facility Award (RP170644). This study was supported by National Institutes of Health grants R01 GM083122 to DN and R35 GM122574 to GW and R01GM112050 to ES.

The authors declare no competing financial interests.

## Author contributions

GF and DN conceived the study and designed experiments. GF and KS performed cryo-ET data collection and analysis. LZ, ED, and YH analyzed the phenotypes of mutants and generated rescued strains. GF and LZ analyzed the LC-MS/MS data. ED and LZ performed sucrose gradient and western blot analyses. NP and LZ performed genotyping analysis of the CLiP mutants. GF and DN wrote the manuscript, and all authors edited the manuscript. The three-dimensional averaged structures of the CA from different strains have been deposited in the Electron Microscopy Data Bank (EMDB) under accession codes EMD-20160 (wild type), EMD-20161 (*fap92*), EMD-20162 (*fap76-1*), EMD-20163 (*fap76-2*), EMD-20164 (*fap81*), EMD-20165 (*fap216*), EMD-20166 (*fap76-1*;*fap81*).

## Supplemental Figure Legends

**Figure S1.**
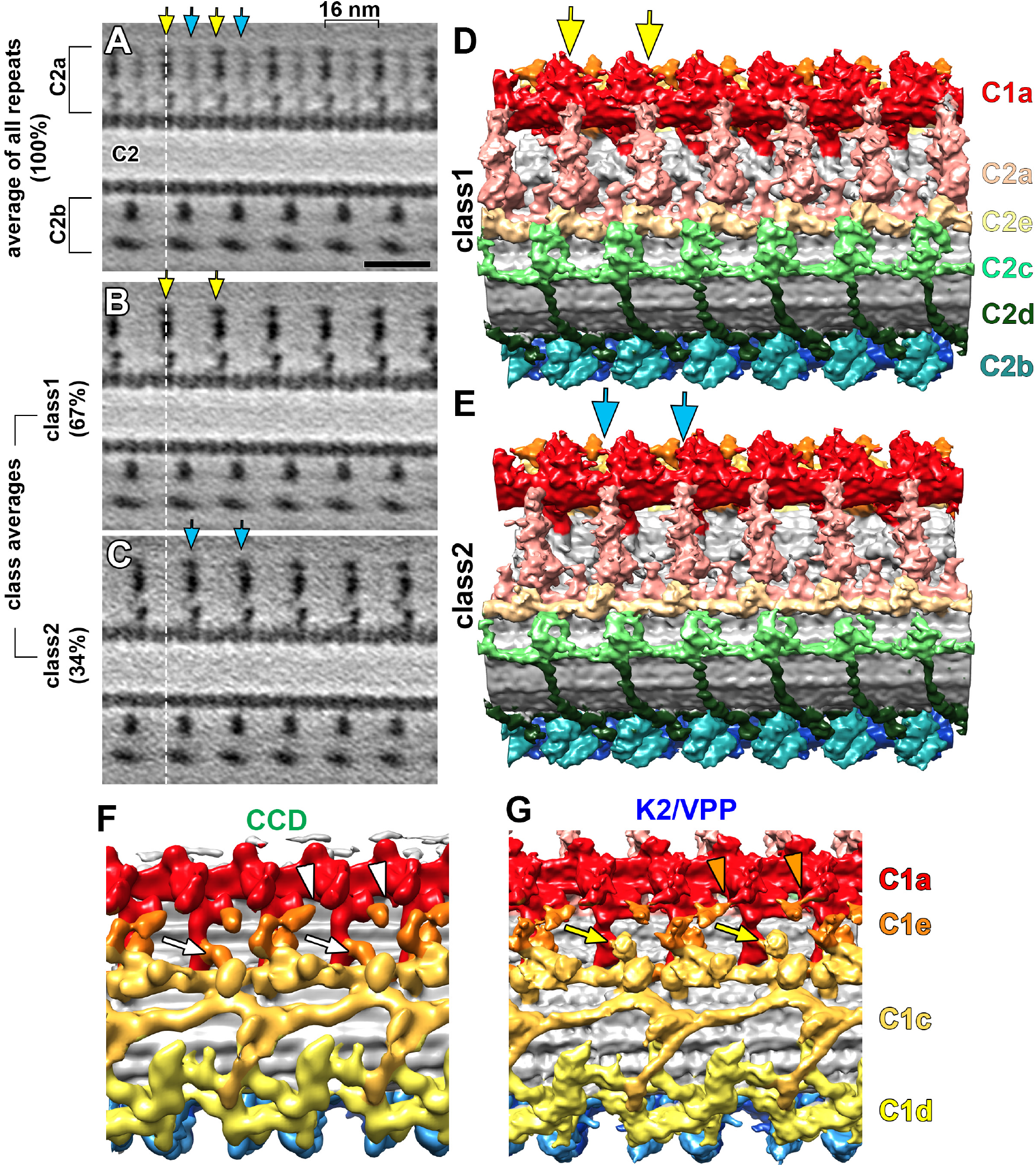
The improved structural resolution reveals two conformational states of the C2a projection and previously unseen structural details of the C1a, C1e and C1c projections. **(A)** Tomographic slice in longitudinal orientation of the averaged *Chlamydomonas* wild-type CA (all repeats, 100%) using advanced hardware for imaging (Krios, K2, VPP), shows strong (yellow arrowheads) and weak (cyan arrowheads) C2a densities suggesting heterogeneity. **(B-E)** A classification analysis revealed two distinct conformational states of the C2a projection, i.e. class 1 (67% of the repeats, yellow arrows in B, D) and class 2 (34% of the repeats, cyan arrows in C, E). The results show that the C2a projection has a 16 nm periodicity, and the dashed line in (A-C) indicates the position of a C2a projection in the class 1 state (B) that shifted 8 nm in the class 2 state (C). **(F** and **G)** The comparison of isosurface renderings between the CCD (F) and K2/VPP (G) CA averages shows that connections between the C1a and C1e projections (orange arrowheads in G) and a peripheral density at the C1c projection (yellow arrows in G) could be visualized in the K2/VPP average but not the CCD data (white arrowheads and arrows in F). Scale bar in A, 20 nm (valid for A-C).

**Figure S2.**
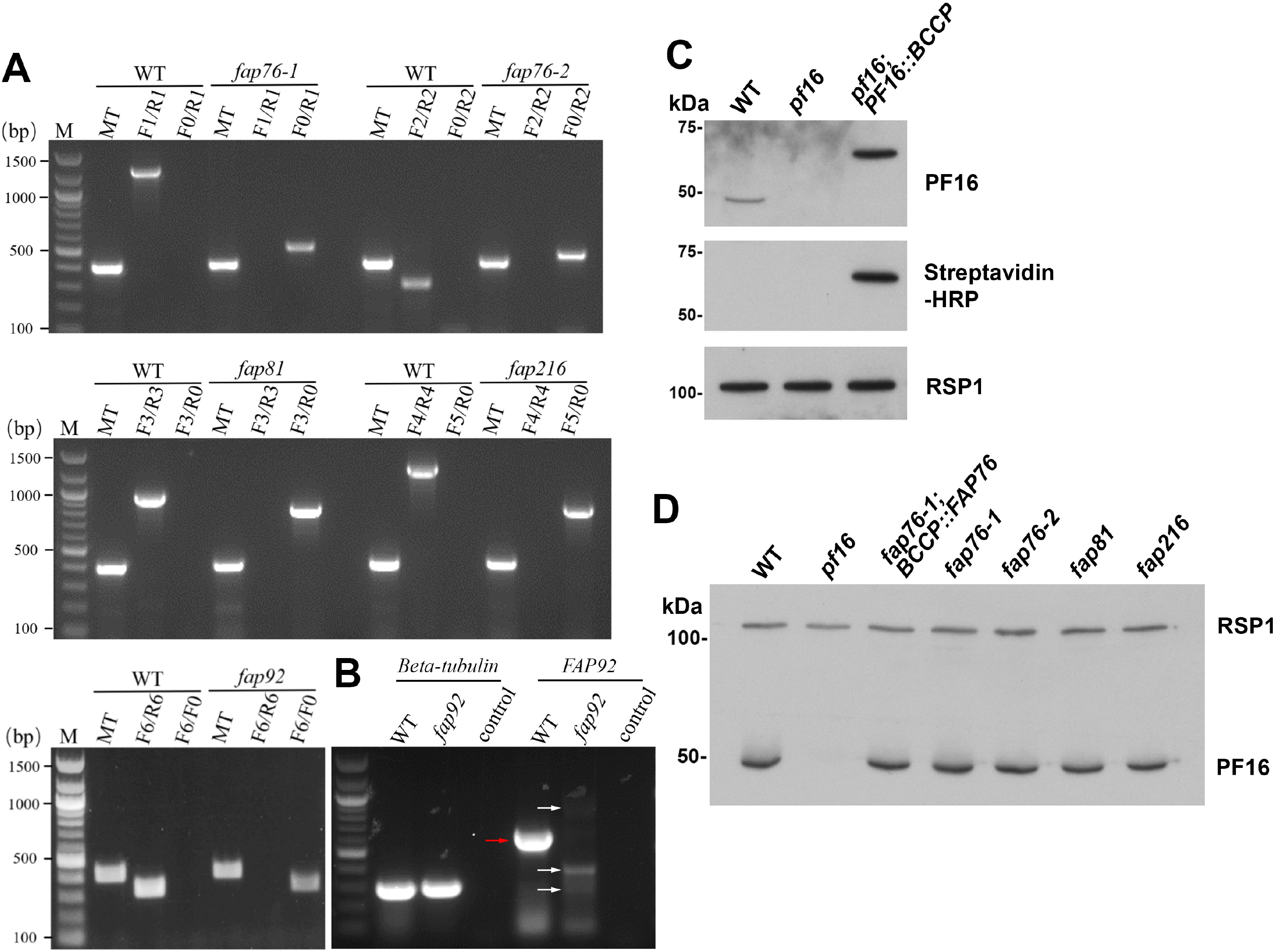
Genotype analyses and western blots probing PF16 in the indicated strains. **(A)** PCR analyses of the *Chlamydomonas fap76-1*, *fap76-2*, *fap81*, *fap92* and *fap216* mutants verified that the corresponding gene was disrupted by the insertion cassette. For each strain, the gene-specific primers could amplify the DNA fragments in wild type, but not in the mutant; in contrast, the cassette fragment could be amplified in the mutant, but not in the wild type. The primers used here are listed in Table S4. **(B)** RT-PCR analyses of the *Chlamydomonas* wild-type and *fap92* mutant revealed that the mRNA of *FAP92* was interrupted in the mutant. Red arrow indicates the right band of *FAP92* cDNA in wild-type confirmed by sequencing and the white arrows indicate the non-specific bands that were not *FAP92* cDNA fragment confirmed by sequencing. *Beta-tubulin* was used as the control gene. **(C)** Western blot analyses using anti-PF16 antibody and streptavidin-HRP show PF16 and BCCP-tagged PF16 in the axonemes of wild type and the rescued strains *pf16;PF16::BCCP*, respectively; the BCCP-tag adds 9 kDa to PF16. In contrast, PF16 was not detected in the *pf16* mutant. The wild-type PF16 co-migrates with a large amount of tubulin and therefore the wild-type PF16 band typically appears of lower intensity than those of the larger PF16-BCCP in the rescued strains. Immuno-labeling of the radial spoke protein 1 (RSP1) was used as loading control. **(D)** Western blot analyses using anti-PF16 antibody showed that PF16 is assembled into the axonemes of the CLiP mutants *fap76-1*, *fap76-2*, *fap81*, and *fap216* and the rescued strain *fap76-1;BCCP::FAP76*. Immuno-labeling of the radial spoke protein 1 (RSP1) was used as loading control.

**Figure S3.**
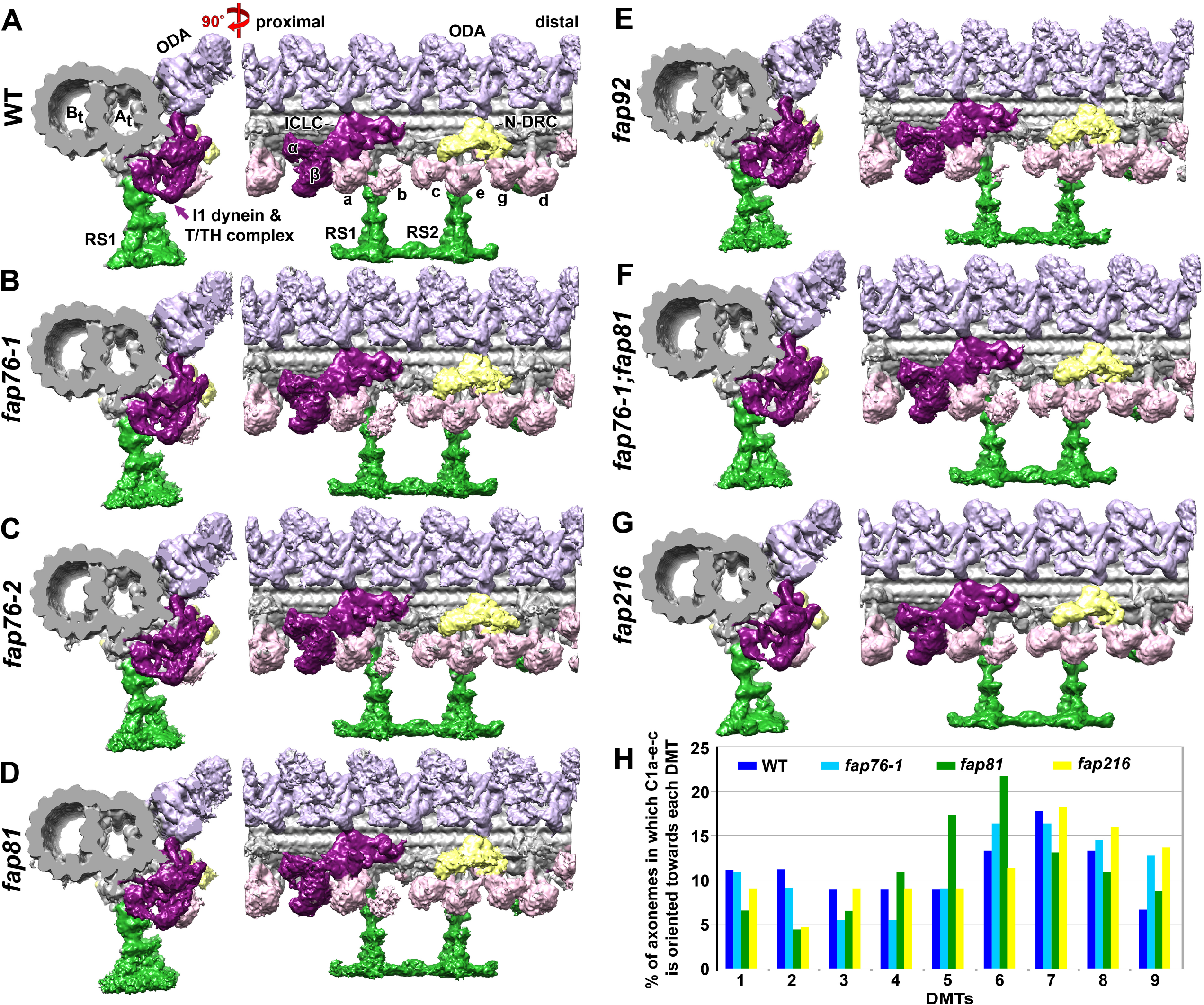
Averages of the 96 nm axonemal repeat of wild type and the indicated mutant strains shows no DMT-associated structural defects in the mutants; also the C1a-e-c supercomplex is preferentially oriented towards DMTs 6-8 in isolated and inactive axonemes. **(A**-**G)** Comparison of the isosurface renderings of the averaged 96 nm axonemal repeats of wild type (A), *fap76-1* (B), *fap76-2* (C), *fap81* (D), *fap92* (E), *fap76-1;fap81* double mutant (F) and *fap216* (G) viewed in cross-sectional (left) and longitudinal (right) orientations shows the same DMT architecture in all strains. Labels: A_t_, A-tubule; B_t_, B-tubule; ODA, outer dynein arm; I1, inner dynein arm I1; α, β, motor domains of alpha and beta heavy chains of I1 dynein; ICLC, intermediate and light chain complex of I1 dynein; a-g, different inner dynein arm isoforms; RS, radial spoke; N-DRC, nexin-dynein regulatory complex; T/TH, tether and tether head. **(H)** Histogram showing the percent of axonemes in which the C1a-e-c supercomplex is oriented toward each of the nine DMTs. The C1a-e-c supercomplex was preferentially orientated towards DMTs 6-8 in wild type and the *fap76-1*, *fap81*, and *fap216* mutants.

**Figure S4.**
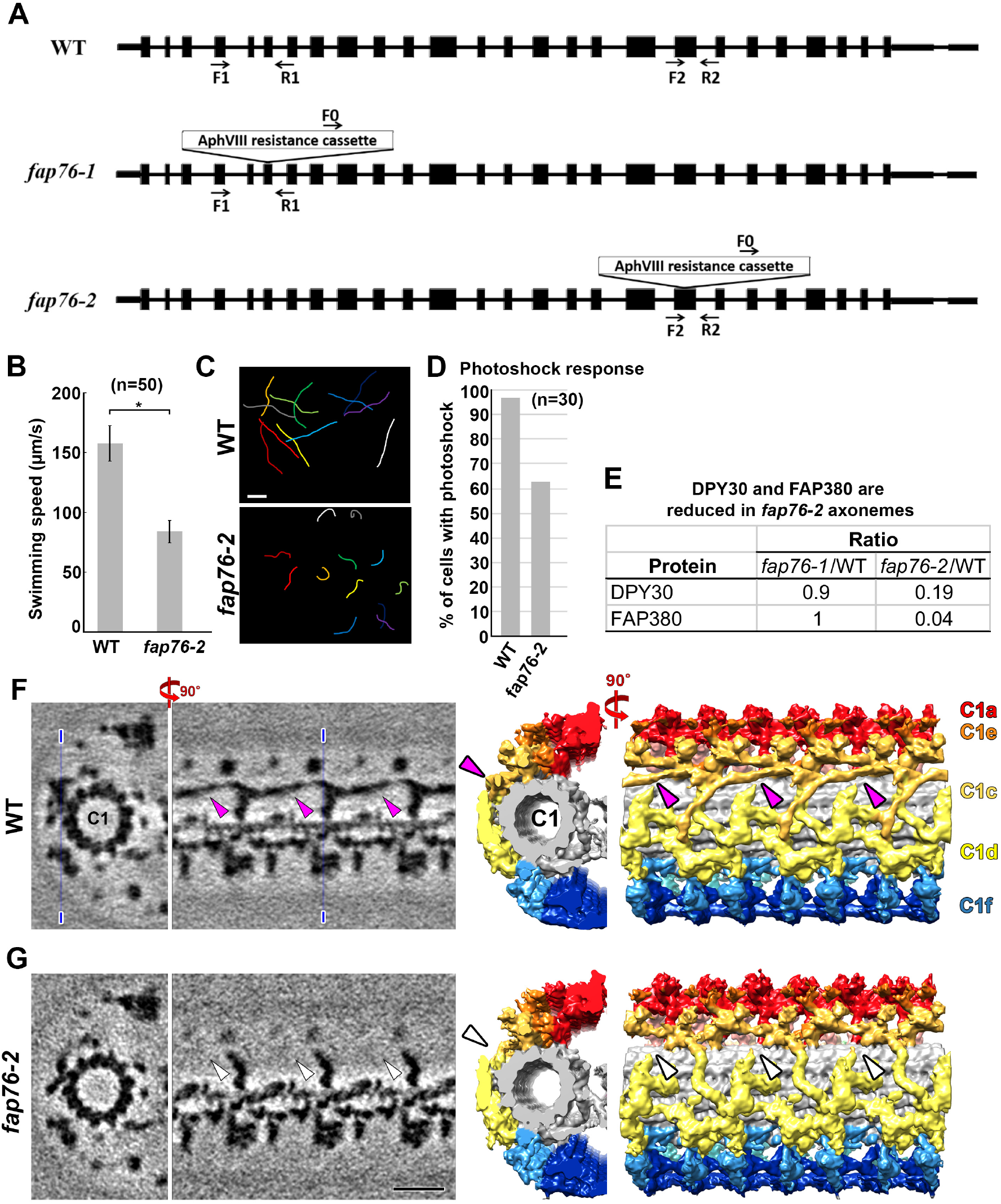
Phenotype and cryo-ET analyses show that the motility and structural defects of *fap76-2* are highly consistent with those observed for *fap76-1*. **(A)** Schematic drawing shows the positions of the cassettes inserted into the *fap76* gene in the *fap76-1* and *fap76-2* mutants. PCR using the indicated primers confirmed the insertion sites (Figure S1 A). **(B-D)** Phenotype analyses show that *fap76-2* cells have slower swimming velocity (B), curving swimming paths (C), and are less likely to undergo the photoshock response (D). **(E)** Mutant/wild-type ratios for two proteins that were greatly reduced in *fap76-2* but remained at wild-type level in *fap76-1*. **(F** and **G)** The comparison of tomographic slices (left) and isosurface renderings (right) between averaged CA repeats of wild type (F) and *fap76-2* (G) revealed the same structural defects in *fap76-2* that were also observed for *fap76-1* (compare to Fig. 4 A-H). The thin blue lines in (F) indicate the location of the tomographic slices viewed in the EM images. Scale bar in C (WT), 50 µm (valid for both WT and *fap76-2* images); in G, 20 nm (valid for all EM images in F and G).

**Figure S5.**
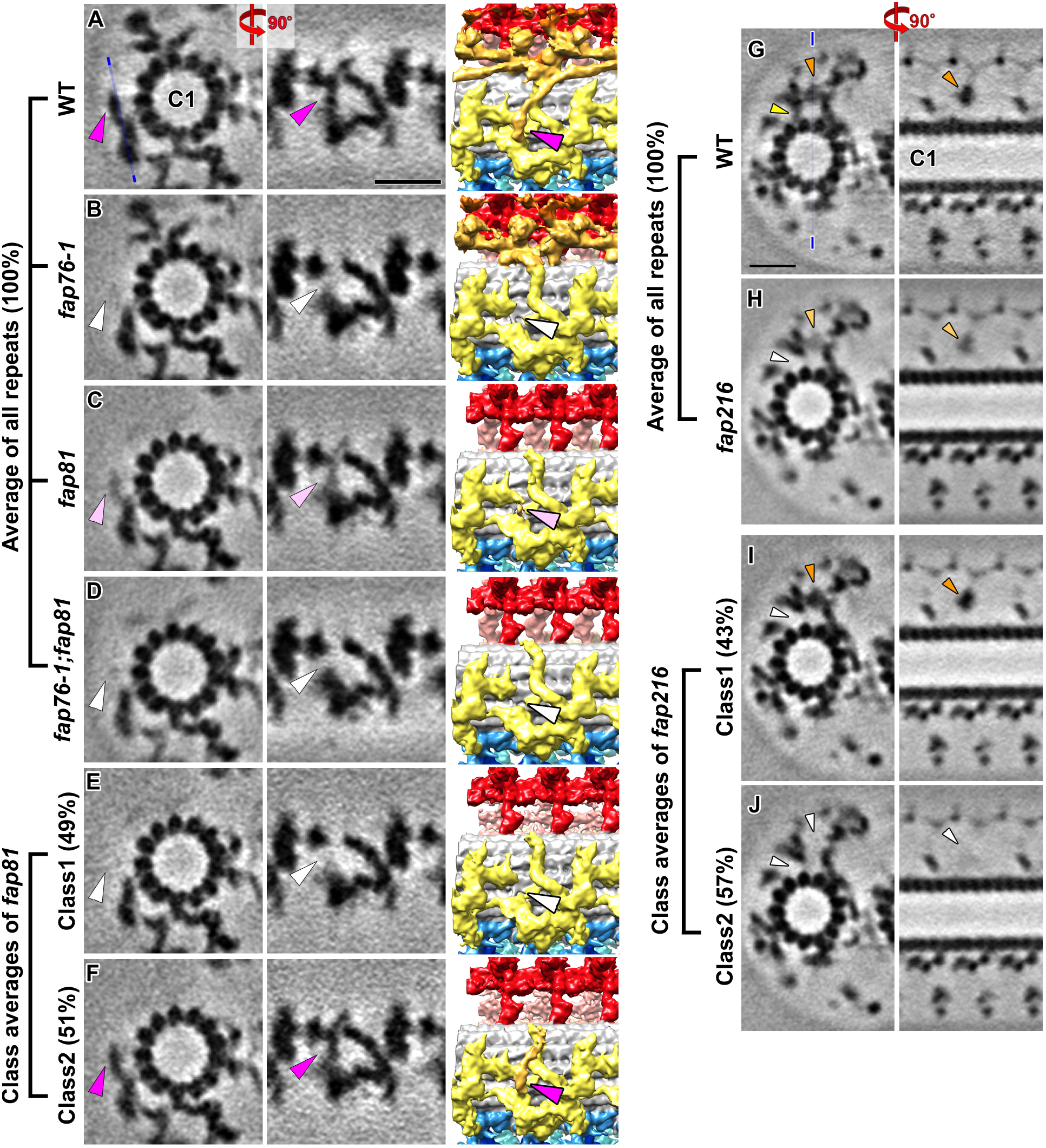
Classification analyses revealed remnant but positionally flexible FAP76 protein in the *fap81* CA and part of the C1e density that bridges between the C1c and C1a projections is reduced in the *fap216* mutant. **(A-D)** The comparisons of tomographic slices (columns 1 and 2) and isosurface renderings (column 3) between the CA averages (all repeats, 100%) of wild type (A), *fap76-1* (B), *fap81* (C) and *fap76-1;fap81* double mutant (D) viewed in cross-sectional (column 1) and longitudinal (columns 2 and 3) orientation, shows that the entire density of wild-type FAP76 (magenta arrowhead in A) is missing in the *fap76-1* and *fap76-1;fap81* double mutant (white arrowheads in B and D), whereas in *fap81* a weak density remains visible where FAP76 usually connects to C1d densities (light magenta arrowhead in C). **(E** and **F)** A classification analysis of the CA repeats in *fap81* focusing on the area where FAP76 interacts with the C1d projection (connection #3) revealed that in half of the CA repeats FAP76 was completely missing (class 1: 49% of the CA repeats), whereas in the other half the FAP76 density was partially visible (class 2: 51% of the CA repeats; magenta arrowhead in F). Note that the FAP76 N-terminal region, which would interact with FAP81 in the wild-type CA, was not observed in the averages, suggesting positional flexibility. **(G** and **H)** The comparison of tomographic slices of averaged CA repeats from all subtomograms of wild type (G) and *fap216* mutant (H) viewed in cross-sectional (left) and longitudinal (right) orientations shows that the FAP216 density that is present in wild type (yellow arrowhead in G) is completely missing in the *fap216* mutant (white arrowhead in H), and a C1e density that bridges the C1c projection to the C1a projection (orange arrowhead in G) is reduced in the mutant (light orange arrowhead in H). The thin blue line in (G) indicates the location for the tomographic slices viewed in longitudinal orientation. **(I** and **J)** Classification analysis focusing on the C1e density that bridges the C1c and C1a projections resulted in two class averages: in class 1 the density was present (43% of the *fap216* CA particles; indicated by orange arrowheads) and in class 2 the density was missing (57% of the *fap216* CA particles; indicated by white arrowheads). Scale bar in A, 20 nm (valid for A-F); in G, 20 nm (valid for G-J).

**Video 1.** Animated 3D visualization of the 32 nm repeat of the *Chlamydomonas* wild-type CA (data collected using K2/VPP). Three averaged repeats spanning a total length of 96 nm are shown in cross-sectional orientation at the beginning of the animation. Nomenclature and color coding of the CA projections are indicated in the animation and follow that of Carbajal-González et al. (2013).

**Video 2.** Summary animation showing the precise locations of FAP76, FAP81, FAP92, and FAP216 within the C1a-e-c supercomplex. The C-terminus of PF16 and N-terminus of FAP76 are also indicated. Nomenclature and color coding of the CA projections are indicated in the animation.

**Video 3.** Collection of videos showing swimming cells of *Chlamydomonas* wide type, the mutants *fap76-1*, *fap76-2*, *fap81*, *fap76-1;fap81* double mutant, *fap92*, *fap216*, and the two rescue strains *fap76-1;BCCP::FAP76* and *fap216;BCCP::FAP216*. The videos are played at 20 frames/s and the cell strain is indicated in the video. Note the uncoordinated ciliary beating in *fap76-1*, *fap81,* and *fap216* mutant cells.

## Footnotes

1 For consistency, we follow the previous nomenclature for CA projections (Carbajal-González et al., 2013).

